# High-throughput analysis of adaptation using barcoded strains of *Saccharomyces cerevisiae*

**DOI:** 10.1101/731349

**Authors:** Vincent J. Fasanello, Ping Liu, Carlos A. Botero, Justin C. Fay

**Affiliations:** Division of Biology and Biomedical Sciences; Washington University School of Medicine in St. Louis; St. Louis, Missouri, USA; Department of Genetics; Washington University in St. Louis; St. Louis, Missouri, USA; Department of Biology; Washington University in St. Louis; St. Louis, Missouri, USA; University of Rochester; Rochester, New York; 14627

## Abstract

**Background:** Experimental evolution of microbes can be used to empirically address a wide range of questions about evolution and is increasingly employed to study complex phenomena ranging from genetic evolution to evolutionary rescue. Regardless of experimental aims, fitness assays are a central component of this type of research, and low-throughput often limits the scope and complexity of experimental evolution studies. We created an experimental evolution system in *Saccharomyces cerevisiae* that utilizes genetic barcoding to overcome this challenge.

**Results:** We first confirm that barcode insertions do not alter fitness and that barcode sequencing can be used to efficiently detect fitness differences via pooled competition-based fitness assays. Next, we examine the effects of ploidy, chemical stress, and population bottleneck size on the evolutionary dynamics and fitness gains (adaptation) in a total of 76 experimentally evolving, asexual populations by conducting 1,216 fitness assays and analyzing 532 longitudinal-evolutionary samples collected from the evolving populations. In our analysis of these data we describe the strengths of this experimental evolution system and explore sources of error in our measurements of fitness and evolutionary dynamics.

**Conclusions:** Our experimental treatments generated distinct fitness effects and evolutionary dynamics, respectively quantified via multiplexed fitness assays and barcode lineage tracking. These findings demonstrate the utility of this new resource for designing and improving high-throughput studies of experimental evolution. The approach described here provides a framework for future studies employing experimental designs that require high-throughput multiplexed fitness measurements.

## Introduction

Experimental evolution in microorganisms such as yeast, bacteria, and viruses has been used to answer evolutionary questions that are experimentally intractable in organisms with longer generation times (de Varigny 1892; Garland and Rose 2009; Kassen 2014; Van den Bergh et al. 2018). A central benefit of most microbial experimental evolution systems is the ability to replicate and repeat evolution as well as the ability to store and compete evolved, intermediate, and ancestral strains (Van den Bergh et al. 2018). Indeed, if there is a single unifying theme to what we have learned from experimental evolution it is that adaptation is universal and often repeatable due to parallel changes down to the molecular level (Burke, Liti, and Long 2014; Kohn and Anderson 2014; Bailey et al. 2017; Graves et al. 2017; Bailey, Guo, and Bataillon 2018). The power of this approach has led to numerous investigations of the effects of population size (Schoustra et al. 2009; A. C. Gerstein et al. 2011; Bailey et al. 2017) and structure (Bell and Gonzalez 2011; Kryazhimskiy, Rice, and Desai 2012; Low-Décarie et al. 2015), mutation rate (Lenski, Sniegowski, and Gerrish 1997; Loewe, Textor, and Scherer 2003; Perfeito et al. 2007; Swings et al. 2017), and various environmental treatments (B. S. Hughes, Cullum, and Bennett 2007; R. Dhar et al. 2011; Riddhiman Dhar et al. 2013; Zhou et al. 2013; Horinouchi et al. 2015; Huang et al. 2018) on fitness and other evolutionary and ecological outcomes.

Measuring adaptation by changes in fitness (i.e., growth rate) is a core requirement of experimental evolution that often limits its implementation. Fitness assays involve direct competition of individuals or populations against one another or a common reference, and traditionally have been carried out using neutral markers scored by plating assays. For example, the long-term experimental evolution conducted by Lenski and colleagues is scored by counting colonies that can ferment arabinose based on colony color (Lenski et al. 1991). Many recent studies use fluorescently marked strains such that the fitness of co-cultured strains can be assayed by flow-cytometry (Gresham et al. 2008; A. C. Gerstein et al. 2011; Selmecki et al. 2015). However, the number of fitness assays, and thereby the resolution of those assays, remain limited, which poses a challenge for large-scale projects. The primary constraint limiting the scale of these studies is that measuring fitness requires replicate assays, often under a variety of conditions. Equally important is the ability to detect small changes in fitness, which is directly related to the number of replicate assays per strain and the noise of the assay itself. Consequently, there is strong motivation to increase the number and sensitivity of fitness assays.

In a barcoding approach, each strain included in the experiment is marked by insertion of a unique DNA sequence (i.e., a barcode) into its genome. Microarrays (Roth et al. 2009) and more recently, direct sequencing of barcodes (Giaever and Nislow 2014), have been used to measure the effects of thousands of single gene deletions on fitness using this approach. Barcodes can also be used to track lineages during experimental evolution and measure fitness improvements via competition assays. Crucially, utilization of barcodes vastly increases the throughput of experimental evolution fitness assays. For example, Blundell, Levy, and colleagues utilized a pool of half a million barcoded yeast strains to detect and track the fate of lineages and to simultaneously measure fitness of thousands of strains relative to one another over a short (<200 generation) time period (Blundell and Levy 2014; Levy et al. 2015). In subsequent work, the fitness of isolates from the evolved population of barcoded strains was measured using short-term competition assays with a common barcoded reference strain (Venkataram et al. 2016; Y. Li et al. 2018). While the above system leverages the throughput of pooled competition assays, it is limited to a small number of evolving populations over short time-scales. However, a similar system could accommodate a large number of evolving populations while retaining the throughput of pooled competition assays if each population were started with a unique barcoded strain(s). Such a system is quite desirable as the number of evolving populations determines the number of treatments, e.g. environments, and the number of replicates.

In this study we describe an experimental evolution system in *Saccharomyces cerevisiae* (hereafter, yeast) that utilizes genetic barcodes to enable high-throughput assessment of adaptation. The system is composed of a library of isogenic, diploid strains that only differ by a unique 20 bp barcode inserted upstream of the *HO* locus. The main advantage of this system is that it enables pooled fitness assays of replicate or differently treated lines marked by unique barcodes. Thus, populations can each be initiated with a single barcode and fitness of the resulting evolved lines can be measured in a single pooled fitness assay. However, the system is also quite flexible, if populations are initiated with multiple barcodes they can be used to track the adaptive dynamics of different lineages that occur during the experimental evolution period. While our system is limited in the number of lineages it can track compared to other studies (Kao and Sherlock 2008; Blundell and Levy 2014; Levy et al. 2015; Selmecki et al. 2015), cross-contamination between populations with different barcodes can be detected and monitored and multiple treatments can be directly compared even if represented by low-relative-fitness lineages.

To demonstrate the capabilities of our system, we began by conducting a series of proof-of-concept fitness assays and subsequently applied what we learned in a short, 25-day (∼250-generation) experimental evolution. To take advantage of the flexibility of the system, the experimental design consisted of 76 populations spanning six different treatments, each initiated with two barcodes per population. We confirm that barcode insertions (1) do not alter the fitness of our source strains, and (2) provide a high-throughput means of measuring fitness differences and evolutionary dynamics for individual lineages from pooled samples obtained at different stages of experimental evolution. We highlight the advantages of multiplexing samples with indexing as well as the importance of limiting molecular contamination between initial sampling and library construction. This system represents a valuable resource for designing and improving high-throughput studies in experimental evolution.

## Materials & Methods

### Strains, media and culture methods

Barcoded yeast strains were constructed using two isogenic haploid derivatives of a strain collected from an Oak tree in Pennsylvania (YPS163) (Sniegowski, Dombrowski, and Fingerman 2002): YJF153 (MATa, *HO*::dsdAMX4) and YJF154 (MATalpha, *HO*::dsdAMX4) (X. C. Li and Fay 2017). To initiate the barcoding process, barcoded kanMX deletion cassettes were amplified from the MoBY plasmid collection (Ho et al. 2009) with primers containing homology to the promoter region (−1,129 to −1,959) of *HO*. This region was selected because without *HO* it is unlikely to be functional. A set of 184 of these barcoded cassettes were selected based on confirmation of correct barcodes by sequencing (Supplemental Table S1 contains entries for barcodes used in this project), then transformed into YJF153 and confirmed by PCR. Barcoded diploid strains were made by mating these barcoded haploids (YJF153) to YJF154, diploids were confirmed by mating-type PCR (Huxley, Green, and Dunbam 1990). The final barcoded yeast strain library consisted of 184 haploid and 184 diploid strains isogenic except for their barcodes; 92 strains from this set were utilized in the proof-of-concept fitness assays and 77 strains from the set were utilized in the 250-generation experiment. The “ancestral reference strain,” discussed below, was arbitrarily selected from amongst the 184 candidate barcodes in the library; the same barcode sequence (1H10) is used for haploids (strain h1H10) and diploids (strain d1H10). Strain construction is a straightforward process that follows accepted protocols; additional strains can be constructed from the MOBY plasmid collection (Ho et al. 2009) as needed. Strains were stored at −80°C as 15% glycerol stocks.

All evolution and fitness assays were conducted in complete medium (CM; 20 g/l dextrose, 1.7 g/l yeast nitrogen base without amino acid and ammonium sulfate, 5.0 g/l ammonium sulfate, 1.3 g/l dropout mix complete without yeast nitrogen base) with or without additional stresses in 96-deep well plates (2.2-ml poly-propylene plates, square well, v-conical bottom; Abgene AB-0932) covered with rayon acrylate breathable membranes (Thermo Scientific, 1256705). Growth plates were incubated at 30°C for 24 hours inside an incubator (VWR, Forced Air Incubator, basic, 120v, 7 cu. ft.) with agitation using a horizontal electromagnetic microplate shaker (Union Scientific LLC, 9779-TC). Saturated (stationary phase) 24-hour culture was diluted (1:1000) into fresh medium at the same time each day to initialize the next round of growth for all evolution and fitness assays.

Starting material for all evolution and fitness assays originated from −80°C freezer stocks of the constructed barcoded yeast strains. Yeast were revived from −80°C freezer stocks via a single round of growth (24 hours, 10 generations) under standard culture conditions (see previous paragraph) for these assays. Yeast samples collected during our experiments were stored both as (1) 15% glycerol stocks at −80°C to maintain viable freezer stocks of yeast populations [i.e., a “frozen time vault”, Van den Bergh et al. 2018] and (2) pelleted samples at −20°C for DNA extraction.

### Experimental design

#### Proof-of-concept fitness assays

The design of this new system for experimental evolution began with a proof-of-concept analysis that allowed us to optimize our methods and to confirm that the fitness of multiple strains in pooled samples could be simultaneously and accurately measured with sequencing-based competition assay methods. This initial step involved measuring the fitness of 91 barcoded yeast strains relative to an ancestral reference strain simultaneously using a sequencing-based fitness assay (Figure 1, A.). Ten replicate pooled fitness assays were conducted under standard culture conditions (see *Strains, media and culture methods*) with the null expectation that all strains should have identical fitness values if the barcode insertion process does not affect fitness. Briefly, 92 strains were revived from −80°C freezer stocks, mixed in equal proportions (i.e., such that each strain comprised 1/92 of the pooled population). The pooled strains were then diluted (1:1000) into fresh medium and grown for two-days, approximately 20-generations, in ten separate wells for the proof-of-concept fitness assays. Samples were obtained from the undiluted initial mixtures (Time 0-hours) and from the final overnight population cultures (Time 48-hours). Fitness was measured by the change in barcode abundance relative to a ‘reference’ strain (d1H10) from 10 replicates for a total of 910 fitness assays (91 focal barcodes x 10 replicates). See *Fitness calculations*, below, for a full description of the competition-based fitness assay methodology and for calculations of fitness from barcode abundance data. DNA was isolated separately for each fitness assay sample using a ZR Fungal/Bacterial DNA Kit (Zymo Research D6005) in individual 2.0 mL screw-cap tubes following the manufacturer’s instructions. Physical cell disruption by bead-beating was carried out in a mixer mill (Retsch, MM 300) at 30 Hz (1800 min^-1^) for ten minutes (1-minute on, 1-minute off, times ten cycles). Following extraction, DNA was amplified with forward/reverse primers containing a 9-12 bp index for multiplex sequencing. PCR products were quantified, pooled and purified to form a single multiplexed library for sequencing. Additional control samples were also included in the library to track barcode cross-contamination. See *Library construction and sequencing*, below, for a detailed description of the library preparation protocol used for these and all other samples.

**Figure 1.**
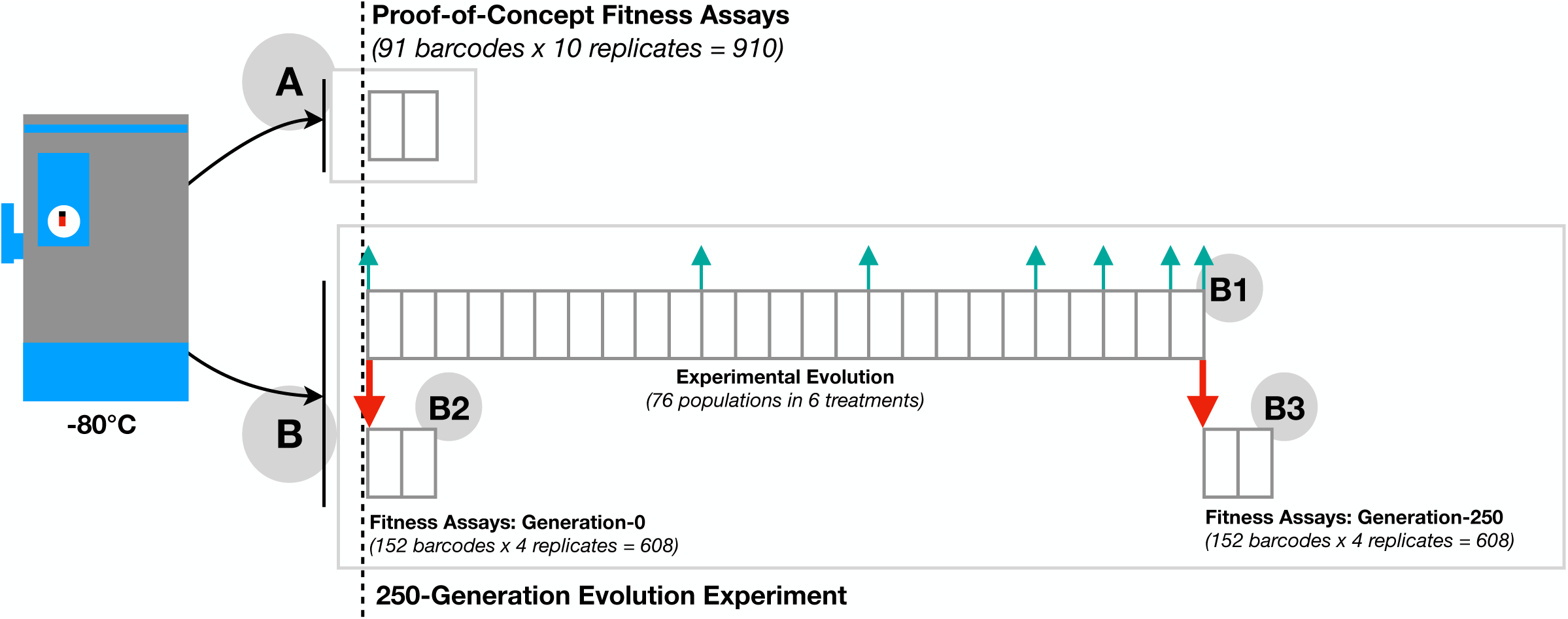
Overview of experimental design. Barcoded yeast strains were used for proof-of-concept fitness assays (A, 2 days) and experimental evolution (B, 25 days) with gray boxes indicating one day (10 generations). The proof-of-concept fitness assay measured fitness by the change in frequency of 91 barcoded strains simultaneously in a pooled population with 10 replicate pools. Experimental evolution (B1) was of 76 populations, each initiated with two barcoded strains per population and experiencing one of six different treatments that had either 11 or 21 replicate populations per treatment. During the 25-day, 250-generation, experimental evolution (B1), evolutionary dynamics samples were collected at generations 0, 100, 150, 200, 220, 240, and 250 (cyan arrows). Fitness of the ancestral (B2) and evolved (B3) strains were quantified via (2-day, 20-generation) fitness assays (red arrows) conducted in several pools (9-23 barcodes per pool), each with four replicates.

#### 250-Generation evolution experiment

After establishing the feasibility of the analysis strategy and optimizing culture, assay, and processing methods, 152 barcoded yeast strains were evolved for 25 days (i.e., ca. 250 generations at 9.97 generations per day – calculated from number of doublings based on optical density data) under different scenarios of selection in a second set of experiments. These 152 strains were paired into groups of two to form 76 populations (wells) for evolution in order to facilitate examination of the evolutionary dynamics within each (see below). Specifically, yeast were evolved by serial dilution in one of six different treatments (Figure 1, B1.; See Supplemental Table S2 for treatment descriptions). Evolutionary treatments involved growth in either complete media (CM), CM with ethanol (8% by volume) or CM with NaCl (0.342 M). Serial transfers were achieved either through standard dilution (1:1000), reduced dilution (1:250), or increased dilution (1:4000). Note that the 1:250 dilution treatment and the 1:4000 dilution treatment are expected to have undergone 200 and 300 generations of evolution, respectively, rather than the 250 generations of evolution expected in the treatments with a standard 1:1000 dilution. This difference in number of generations is caused by variation in the number of possible doubling events when different numbers of cells are added to media with the same (limiting) amount of glucose. Additionally, haploid and diploid yeast were evolved under standard culture conditions (see *Strains, media and culture methods*) to assess the effects of ploidy. To initialize this evolution experiment, yeast were revived from −80°C freezer stocks. Yeast strain pairs slated to evolve in competition were then mixed in equal proportions and diluted in fresh medium, according to the experimental design, and allowed to evolve for 25 days.

Yeast samples were obtained from the initial undiluted mixtures and on day 25 to use as the starting material for the Generation-0 and the Generation-250 fitness assays. These end-point assays of relative fitness were developed to characterize fitness outcomes in the 250-generation experiment (See *End-point assay of relative fitness change*). Additional samples were obtained at regular intervals to characterize evolutionary dynamics (See *Evolutionary dynamics*). These two sets of assays are described in detail in the two following sections.

*Evolutionary Dynamics:* Relative proportions of strain pairs evolved in competition (within the same well) were quantified at generations 0, 100, 150, 200, 220, 240, and 250 (Days 0, 10, 15, 20, 22, 24 and 25). Yeast samples from each timepoint were pooled for DNA extraction such that there was no barcode overlap within pools. DNA was subsequently extracted (see the *Proof-of-concept fitness assays* section for details). Libraries were constructed for sequencing as described in the *Library construction and sequencing* section. From these data, the relative proportions of strains within each pair were measured at each time-point. Using this information, the time-point (t-max) and magnitude (m-max) of the maximum change in relative abundance in comparison to the starting conditions, the time-point (t-max-rate) and magnitude (m-max-rate) of the maximum rate of change between adjacent time-points, the time-point (t-max-diff) and maximum difference (m-max-diff) in barcode proportions, and the total cumulative change in barcode relative abundance were calculated for each two-barcode population. Barcodes approaching fixation (hereafter referred to as “fixed”) were also noted and were defined as cases in which a single barcode in the pair obtained (and maintained) a proportion of 0.95 or greater by (through) generation 250 of the evolution experiment.

*End-point assays of relative fitness:* This second type of assay was designed to assess the fitness of each focal strain relative to a static reference using a barcoded competition-based assay. Using a static reference enabled us to quantify the change in fitness for each strain (relative to the reference) between the start and end of an experimental evolution. In this case, the assays involved quantifying barcode fitness using pooled samples from generation-0 (Figure 1, B2.) and separately pooled samples from generation-250 (Figure 1, B3.) from the 250-generation evolution experiment (Figure 1, B1.). Briefly, Generation-0 and Generation-250 yeast strains were revived and samples from each time point were pooled, separately, such that there was no overlap in barcoded yeast strain identity within each pool. Pools contained 8-22 unique barcodes. Pools were independently combined in equal proportion (50% pooled-yeast : 50% ancestral reference) with an ancestral reference strain (re: an ‘unevolved’ barcoded yeast strain), then diluted into fresh medium to initialize the fitness assays. Four replicate fitness assays were conducted for each pool of generation-0 and generation-250 yeast. Fitness assays employed a standard 1:1000 transfer dilution across all samples (see *Strains, media and culture methods*). Samples evolved in CM plus additional stresses were assayed in the same media type in which they were evolved. Diploids were competed against a diploid reference strain (strain ID: d1H10), while haploids were competed against the haploid version of this same reference (strain ID: h1H10). In these assays, yeast samples for DNA extraction were obtained at hour-0 from the undiluted initial mixtures (fitness assay starting material), and, 20 generations later, at hour-48 from the final overnight population cultures (fitness assay end) to calculate fitness. See *Fitness Calculations*, below, for a full description of fitness assay methodology and for calculations of fitness from fitness assay barcode abundance data. Libraries were constructed for sequencing as described in the *Library construction and sequencing* section, below.

#### Library construction and sequencing

Barcode sequencing libraries for the proof-of-concept fitness assays and all samples from both components of the 250-generation evolution experiment were constructed by amplification of MoBY barcodes (Ho et al. 2009) with primers containing Ion Torrent adaptors with indexes assigned to distinguish samples from one another (Supplemental Table S3).

PCR products for library construction were generated using 25 cycles and were subsequently quantified with a Qubit 3.0 Fluorometer, high sensitivity assay kit. These products were then combined at equimolar concentrations and purified using a Zymo DNA Clean & Concentrator kit (Zymo Research D4014) to create a single library for sequencing. The DNA extraction and PCR steps were repeated for samples that did not attain sufficient DNA concentrations (no samples were exposed to > 25 rounds of PCR). We note that PCR jackpotting can be an issue in DNA library construction and that it can add noise to measurements of barcode frequency. To minimize the influence of this phenomenon on our results, we used a large amount of template DNA (Cha and Thilly 1993), and conducted all fitness assays in four replicates. Additionally, we avoided misclassification of barcodes by using high fidelity Taq Plus DNA Polymerase (Lambda Biotech, Taq Plus Master mix Red), and MOBY barcode sequences that differ from one another by at least six substitutions.

DNA libraries were sequenced using an Ion Torrent sequencer (Ion Proton System, Ion Torrent) at the Genomics Core Facility at Saint Louis University with a customized parameter to assess polyclonality after 31bp (the start position of the forward Ion Torrent adapter index sequence). A single sequencing run was used for each pooled library (library 1 – proof-of-concept fitness assays, library 2 – evolution experiment: evolutionary dynamics samples, Generation-0 fitness assays, and Generation-250 fitness assays). An additional library was constructed and sequenced for a set of samples from library 2 with elevated barcode cross-contamination rates.

### Sequence data processing & calculations

#### Sequence datasets

Sequence data in FASTQ format were parsed and demultiplexed using custom scripts in R (see the *Availability of data and materials* statement, below). A total of 142,243,245 raw reads that matched the forward Ion Torrent adapter indices included in our experiment (omitting reads that matched no forward adapter, polyclonal reads, low quality reads, and adapter dimer reads) were recovered across the sequenced libraries. 104,365,740 reads (73.4%) were retained for analysis that perfectly matched a forward sequencing adapter index (9-12 bp), reverse sequencing adapter index (9-12 bp) pair, and a MOBY genetic barcode (20 bp) included in the full experimental design.

#### Barcode cross-contamination rate

The barcoded yeast experimental evolution system has an innate ability to detect and track barcode cross-contamination that could arise during evolution, over the course of short-term fitness assays, or during DNA library preparation. Barcode cross-contamination rate was defined as the mean number of counts mapping to any given barcoded yeast strain included in the full experimental design (sequenced library), but not expected to be present in the given sample (pair of forward and reverse IonTorrent adapter Indices). Barcode cross-contamination rates were calculated separately for each unique forward-reverse index pair included in each sequencing library and represent the amount of noise an average contaminating barcode strain contributes to each sample. The barcode cross-contamination rate of sample (primer pair) j, B_j_, was measured as,

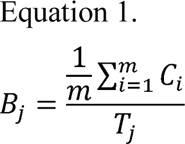

where *m* is the number of barcoded strains that could potentially contribute to barcode cross-contamination in experiment *j* (i.e., the number of unique strain IDs included in the full library, but not expected in sample j), *C_i_* is the number of barcoded yeast strain counts recovered for contaminating barcode *i*, and *T_j_* is the total number of counts recovered in sample *j* across all barcoded yeast strains included in the full experimental design.

Contamination rate summaries are reported separately for the proof-of-concept fitness assays and the evolution experiment (the latter containing the evolutionary dynamics, generation-0 fitness assay and generation-250 fitness assays samples). For the subset of evolution experiment samples that were sequenced in two separate libraries, only the replicate with a lower contamination rate was retained for contamination rate summary reporting and all downstream analyses. In all statistical analyses reported below, contamination rate is initially included as a potential predictor and subsequently removed from the model if its effect was found to be non-significant.

#### Fitness calculations

Fitness was evaluated via sequencing-based assays that involved competing barcoded yeast strains against a common ancestral reference strain for 48 hours (two overnight cultures; 20 generations). Reported fitness values in all fitness assays are relative to the same ancestral reference strain (strain ID: d1H10 for diploids; strain ID: h1H10 for haploids). The relative Malthusian fitness of barcoded yeast strain *i* at generation *gn, m_i gn_*, was measured as,

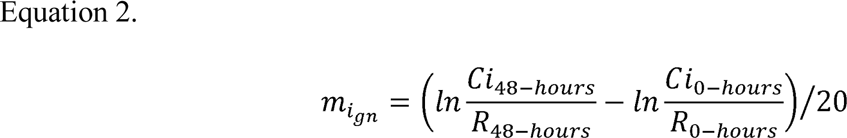

where C*i* and *R* refer to barcode counts for the focal barcode and reference barcode at fitness assay time 0-hours (initial mixtures; fitness assay initial measurement) and time 48-hours (final overnight cultures; fitness assay final measurement), and *20* is the number of generations over 48 hours (two overnight cultures at 9.97 generations each – calculated from number of doublings based on optical density data) (Hartl and Clark 1997; Chevin 2011). We use the standard equation, m=ln(w) to convert Malthusian fitness values to Wrightian fitness values (Orr 2009; Wu et al. 2013; Passagem-Santos and Perfeito 2018). Herafter, fitness, denoted by a *w* will refer to Wrightian fitness. The change in fitness of strain *i* between Day 0 and Day 25 in for our 250-generation evolution experiment, Δ*w_i_*, we therefore computed as,

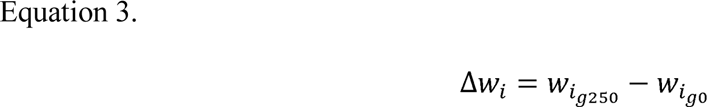

where *w_i g0_* and *w_i g250_* are the strain’s fitness relative to the ancestral reference at generation 0 and 250 as measured from Equation 2. Thus, we assumed no frequency dependent selection.

### Statistical analysis

#### Analysis & visualization tools

All Analyses and statistical test were conducted in R version 3.5.2 (R Core Team 2012). Data processing uses primarily base-R functionality; the plyr package (Hadley Wickham 2011) is used in some cases for data frame manipulation. All Statistical models with linear mixed effects were generated using the lme4 (Bates et al. 2015), and lmerTest (Kuznetsova, Brockhoff, and Christensen 2017) packages. Weighted t-tests were assessed with the weights package (Pasek 2018). Power analyses were conducted using the pwr package (Champely 2018). Finally, figures and tables were generated with the ggplot2 (Wikham 2009) and sjPlot (Ludecke 2019) packages, respectively; multi-panel figures were built using the gridExtra package (Auguie 2017). Raw p-values are reported unless otherwise noted; some tables included in the supplement report raw and corrected p-values (i.e., false discovery rate) for cases where only corrected values are presented in the main text.

#### Reads

Analysis of multiplex barcode sequencing data requires careful consideration of sample size. Because count data are ultimately handled as relative frequencies (proportions), it was necessary to consider underlying sample size or “confidence” in each piece of data within the full dataset for all calculations and analyses. That is, entries with more reads were explicitly assumed to contribute more to summary calculations and statistical models. Variation in sample size was thus controlled for by weighting all calculations by the read sample size and by including such weights in downstream statistical models. This sample size metric considers both the total counts recovered for a multiplexed sample (unique forward and reverse index sequence adapter pair) and the number of counts recovered for the focal barcoded yeast strain(s) in that sample. In the analyses presented here, read calculations utilize harmonic means rather than arithmetic means; the harmonic mean is sensitive to small values and therefore does not overinflate the read values for any variables calculated from a combination of high-and low-read data sources.

#### Proof-of-concept fitness assays

Fitness differences among 91 constructed barcoded yeast strains used in the proof-of-concept fitness assays were assessed using a weighted linear model with mean-corrected fitness as the response variable and yeast strain ID as the predictor variable (mean-corrected fitness ∼ strain ID; family = gaussian). Barcode cross-contamination is not included in this model as all fitness assay samples contained all 91 focal barcodes; barcode cross-contamination is reported from a separate set of 1-barcode and 2-barcode samples for the Proof-of-concept fitness assays.

#### Power Analysis

Power analyses were conducted to calculate the sensitivity of the system in measuring treatment-level fitness effects, fitness changes for individual barcoded yeast strains, and initial fitness variation among individual strains. Power was calculated using sample size and fitness effect as variables of interest with a fixed value for error in fitness measurements. Error in fitness was estimated from the root mean squared error among replicates from the Proof-of-Concept fitness assays as well as from replicate measures of fitness change for individual barcodes from the 250-generation experiment. Power analyses were conducted using the pwr package in R; the pwr.f2.test function was used to calculate power to discern treatment-level differences in fitness and the pwr.t.test function was used to calculate power to detect individual barcode fitness changes and initial fitness differences among individual barcodes.

#### Multiplex barcode sequencing and cross-contamination

Barcode cross-contamination rate was assessed (Equation 1) and summarized, separately for the proof-of-concept fitness assays and the 250-generation experiment. The effects of barcode cross-contamination rate on the change in fitness and the error among replicate fitness measures for each barcode were assessed by including barcode cross-contamination as a predictor in two sets of linear mixed-effects models with either change in fitness over 250 generations of experimental evolution or standard error in fitness change over 250 generations (across four replicate measures for each barcode) as the response variable. Additionally, a weighted 1-tailed t-test was utilized to assess whether barcode cross-contamination rates for samples collected in the 250-generation experiment decreased after re-extracting DNA and resequencing. See the following sections for linear model and linear mixed-effects model specifications.

#### Fitness change in 250 generations of experimental evolution

Individual barcoded yeast strains that exhibited a change in fitness over the 250-generation fitness experiment were identified using a weighted linear model with change in fitness as the response variable and a strain identifier (treatment plus yeast strain Barcode ID) as the predictor variable (fitness change ∼ strain ID; family = gaussian).

Next, the effects of evolutionary treatment (medium type, ploidy, and transfer dilution) on change in fitness over 250 generations of experimental evolution was assessed using a weighted linear mixed-effects model with cross-contamination rates as a covariate. A random effect of strain identifier (treatment plus yeast strain barcode ID) was placed on the model intercept (Fitness change ∼ treatment + cross-contamination + (1|Strain identifier); family = gaussian). A second weighted linear model was fit to assess potential sources of error in the measurement of fitness in this system. Here, a model was constructed using a comprehensive set of factors that could have impacted the standard error in fitness measurement among replicates. These included: treatment, cross-contamination rate, magnitude fitness change, median count (median number of counts returned for a focal barcode across all sampling points that contribute to the fitness change calculation), and median proportion (median proportion of the focal barcode in its mixed 2-barcode well across all sampling points); (SE-Fitness change ∼ treatment + cross-contamination + fitness change + median count + median proportion; family = gaussian).

#### Evolutionary dynamics

For all analyses of evolutionary dynamics, only the first barcoded yeast strain from each pair was included to ensure that data were statistically independent. A set of weighted linear mixed effects models, with treatment as the predictor variable and either t-max, m-max, t-max-rate, m-max-rate, t-max-diff, m-max-diff, or total cumulative change in barcoded relative abundance across all time-points as the dependent variable, were employed to assess treatment differences in evolutionary dynamics. Full models included initial barcode abundance as a predictor term because initial barcode abundance could impact subsequent dynamics (e.g., (t-max ∼ Treatment + Initial barcode abundance; family = gaussian)). Initial barcode abundance was subsequently removed from models when deemed nonsignificant, resulting in removal from all but one model (t-max-diff). Barcode fixation rate was not assessed statistically due to the small number of fixation events observed (n=7/76 populations).

### Availability of data and materials

The dataset supporting the conclusions of this article is available in the NCBI Sequence Read Archive (SRA) repository, BioProject Number PRJNA555990, https://www.ncbi.nlm.nih.gob.bioproject/PRJNA555990 (Fasanello et al. 2019). Data formatted for analysis, intermediate data frames, as well as the custom R scripts utilized for all data processing, statistical analysis, and figure generation are available from GitHub (github.com/VinceFasanello/MM_Code_Supplement); a static version of the repository is available from Zenodo (Fasanello 2020). A readme file is available in the repository with the instructions necessary to reproduce the analyses and to confirm the results presented in this article. Supplementary figures, tables, and files referenced throughout the main text are available as “Supplemental Files”. Strains are available from the Justin C. Fay Lab at The University of Rochester; contact James Miller (e).

## Results

### Proof-of-concept fitness assays

We constructed 92 diploid yeast strains, all genetically identical except for a unique 20 bp barcode inserted upstream of the deleted *HO* gene. Proof-of-Concept fitness assays were then used to measure any fitness differences among these strains, to estimate our power to detect small fitness differences, and to assess our ability to measure fitness using multiplexed barcode sequencing. We assayed fitness simultaneously for our pool of 91 barcodes by measuring changes in barcode abundance relative to a ‘reference’ strain (barcode ID: d1H10) over a two-day period of approximately 20 generations.

A few of the barcoded strains showed significant differences in fitness (3/91 at a 5% false discovery rate, FDR; 1/91 at a 1% FDR) (Figure 2, Supplemental Table S4), possibly due to mutations arising during the barcoding process. The root mean squared error (rMSE) among replicated measures of fitness was 1.76e-2, indicating good power (80%) to detect fitness differences of 1.2% between strains with four replicates at a nominal significance level of P =0.05 (Figure 3a-green lines, Supplemental Table S5). We subsequently used these data to estimate power to detect treatment effects with end-point fitness assays in our proposed experimental design for the 250-generation experiment. With a modest number of barcoded yeast strains per treatment (n=22), we estimate high power (99.8%) to detect 1% fitness differences between treatments at a nominal significance level of P = 0.05 with four replicate end-point fitness assays (Figure 3b - green lines, Supplemental Table S6). With these promising results, we proceeded to conduct a more comprehensive test of the utility of barcoded strains in a practical experimental evolution context (i.e., the utility to evaluate the effects of treatment on fitness outcomes and evolutionary dynamics).

**Figure 2.**
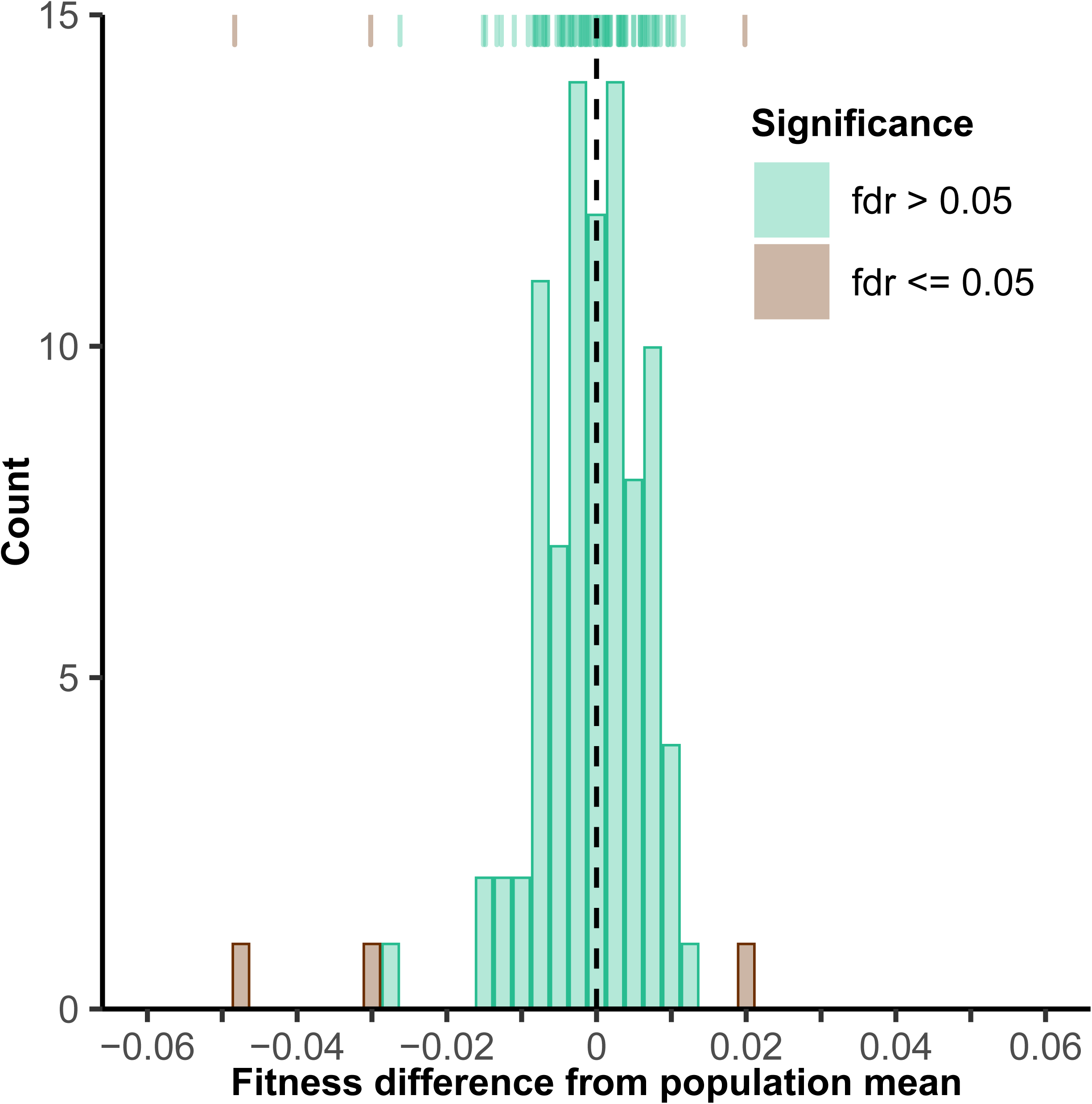
Histogram of fitness from 91 barcoded yeast strains. Fitness is the deviation from the population mean and was quantified by competition against a common reference strain via Proof-of-concept fitness assay. Orange and green bars indicate yeast strains with fitness values significantly and not significantly different from the population mean at a false discovery rate of 5%, respectively.

**Figure 3.**
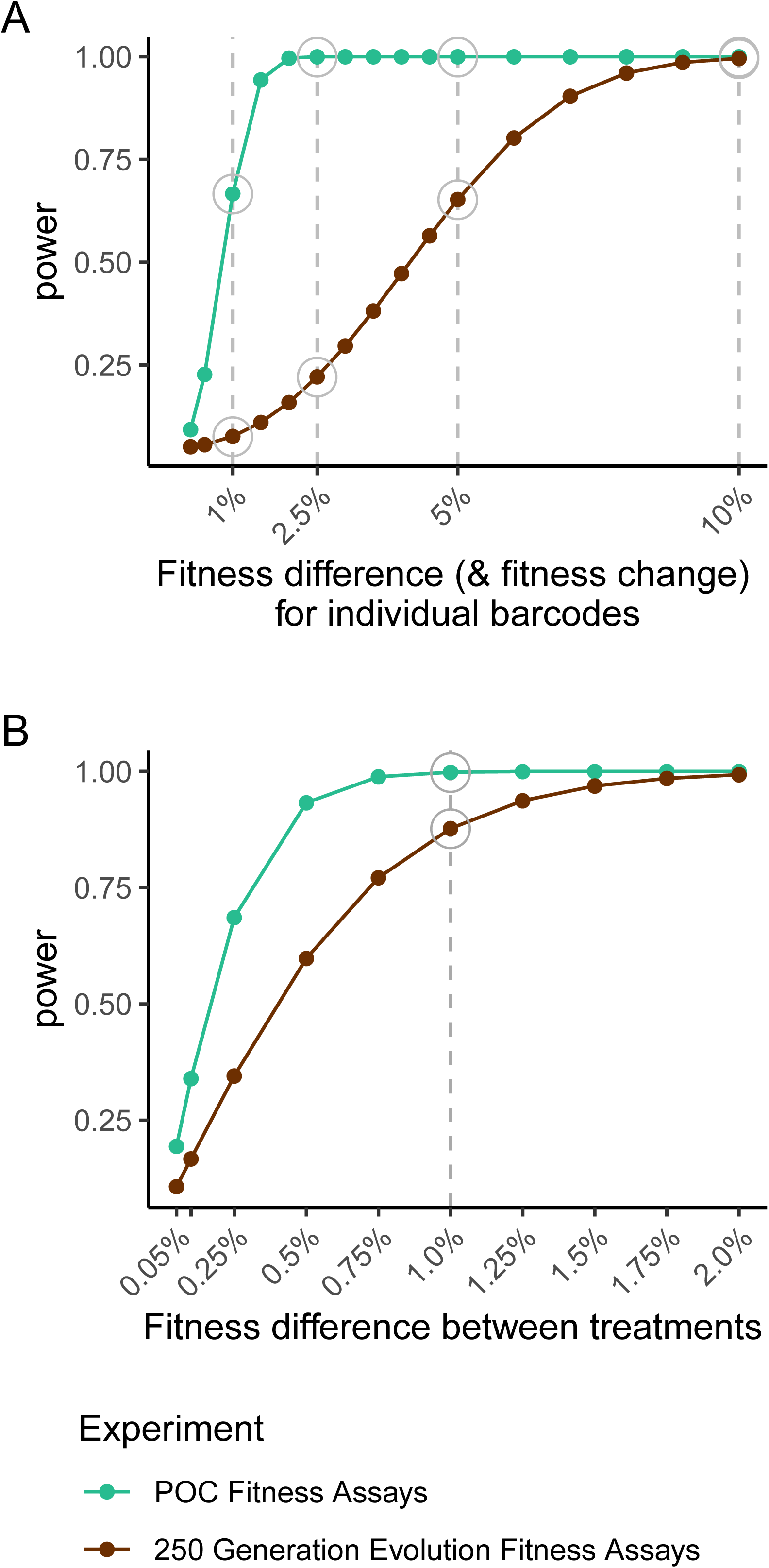
Power surfaces for POC fitness assays data (green) and fitness change data from the 250-generation experiment (orange). Power to detect individual barcode fitness deviation from population mean calculated using error in the POC fitness assay data (A, green) and power to detect individual barcode fitness change using error in the 250-generation experiment (A, orange); both employ four replicate measures of fitness. Power to detect treatment-level fitness differences for two treatments with 22 barcodes in each as estimated using error in the POC fitness assay data (B, green) and the 250-generation experiment (B, orange).

### Experimental evolution

To empirically evaluate the strengths of this barcoded-strain system for studies of experimental evolution, we conducted a 25-day evolution experiment, which is approximately equal to 250 generations of evolution. Our design included 76 populations, each initiated with two barcodes per well. We used two barcodes per well to monitor evolutionary dynamics (Kao and Sherlock 2008; Selmecki et al. 2015) while minimizing the chance of barcode loss (Blundell and Levy 2014). Samples spanned six treatments, which varied in yeast strain ploidy, growth medium, and daily transfer dilution (Supplemental Table S2). To measure any changes in fitness we competed the evolved generation-250 strains or their generation-0 ancestors against a common reference strain over a two-day period of approximately 20 generations. We also measured the change in barcode frequency of barcoded strains within each evolving population (barcode pair) from longitudinal-evolutionary dynamics samples collected at 7 timepoints across the evolution. From these samples we quantified the magnitude and timing of changes in barcode frequency, which should be influenced by changes in the fitness of the evolving barcoded strains present within each population.

#### Multiplex barcode sequencing

We utilized a two-step multiplexed design to obtain high throughput estimates of fitness based on barcode sequencing in our fitness assays. In the first step we leveraged the strain-identifying barcodes by pooling multiple strains together and simultaneously competing them against an ancestral reference strain to estimate relative fitness. In our second multiplexing step we PCR-amplified each fitness assay sample with unique forward and reverse indexed sequencing adaptors. This latter step enabled us to assign sequencing reads to the appropriate fitness assay after sequencing many samples in concert as a single library. We constructed one library for each experiment (Library 1 – Proof-of-Concept fitness assays; Library 2 – 250-generation evolution experiment evolutionary dynamics and fitness assays samples).

From 84,776,627 demultiplexed reads, we obtained a median of 2,961 reads per barcode in each sample. However, we also detected barcode cross-contamination, defined as barcodes with sample indexes that shouldn’t exist in our sequenced libraries. The average rate of barcode cross-contamination per sample was low, 0.04%, and consistently present in nearly all of our samples (Supplemental Figure S1 A, B). Contamination could occur during culturing of yeast strains, liquid handling during preparation of libraries (Lenski et al. 1991; Van den Bergh et al. 2018), or potentially via index switching during sequencing procedures (Illumina 2017; Sinha et al. 2017; Costello et al. 2018). However, the low but uniform rate of cross contamination we observe is more consistent with library preparation and sequencing than yeast contamination. This is further evidenced by the fact that no growth was observed in culture blanks during any fitness assays nor during the 25-day experimental evolution experiment.

To test whether the barcode cross-contamination rate depends on liquid handling during library preparation, we reprocessed a subset of samples with high cross contamination rates using identical starting material (multiple yeast sample aliquots were created at the time of sample collection). Cross contamination rates decreased significantly in these reprocessed samples (t = 22.3, df = 65, p < 10e-6), but a low level of background contamination remained (Original Samples: mean = 0.75%; reprocessed samples: mean = 0.05%) (Supplemental Figure S2).

In our analyses, the presence of low abundance barcode cross-contamination was removed from all samples in which a barcode is not expected to occur. However, this doesn’t eliminate contamination in samples where a barcode is expected to occur. In such cases, error in estimates of barcode frequency is highest when a barcode’s frequency is low and approaches the cross-contamination rate. The effects of contamination rate and other system-specific covariates on experimental outcomes are presented in detail in the next section.

#### End-point fitness assays

We assessed fitness in the 152 evolved strains (evolution experiment generation-250 strains) as well as their ancestors (evolution experiment generation-0 strains). For each strain and timepoint, fitness was measured in comparison to a common ancestral reference strain via a two-day competition-based fitness assay in the same media type under which the strain had been evolved. The root mean squared error (rMSE) among replicated measures of fitness was 4.31e-2 for these assays. This is slightly higher than our proof-of-concept fitness assays (rMSE = 1.76e-2) but is still expected to provide reasonable power to detect fitness changes for individual barcodes (80% power to detect fitness changes of 2.99%) and fitness differences among treatments (80% power to detect treatment differences of 0.81%) (Figure 3 - orange lines, Supplemental Table S7, Supplemental Table S8). Among a number of covariates tested we found that only the cross-contamination rate (P = 0.01) and magnitude fitness change (P = 6.03e-4) were associated with error among replicates (Supplemental Table S9).

Twenty three percent (35/152) of the strains showed significant increases in fitness between generation 0 and generation 250 at a 1% false discovery rate, FDR (Figure 4, Supplemental Table S10). Within this subset, the average fitness increase was 6.80% with a range of 2.75% to 23.5%. Relatively few strains with significant increases in fitness were found in the CM treatment (3/42; 7.14%) and ethanol stress treatment (2/22; 9.09%). A higher proportion of strains exhibited significant increases in fitness in the lower 1:250 dilution treatment (9/22; 40.9%), the salt stress treatment (11/22; 50.0%), and the haploid treatment (10/22; 45.5%). No strains in the 1:4000 dilution treatment (0/22) exhibited increases in fitness. No strains (0/152) exhibited a significant decrease in fitness.

**Figure 4.**
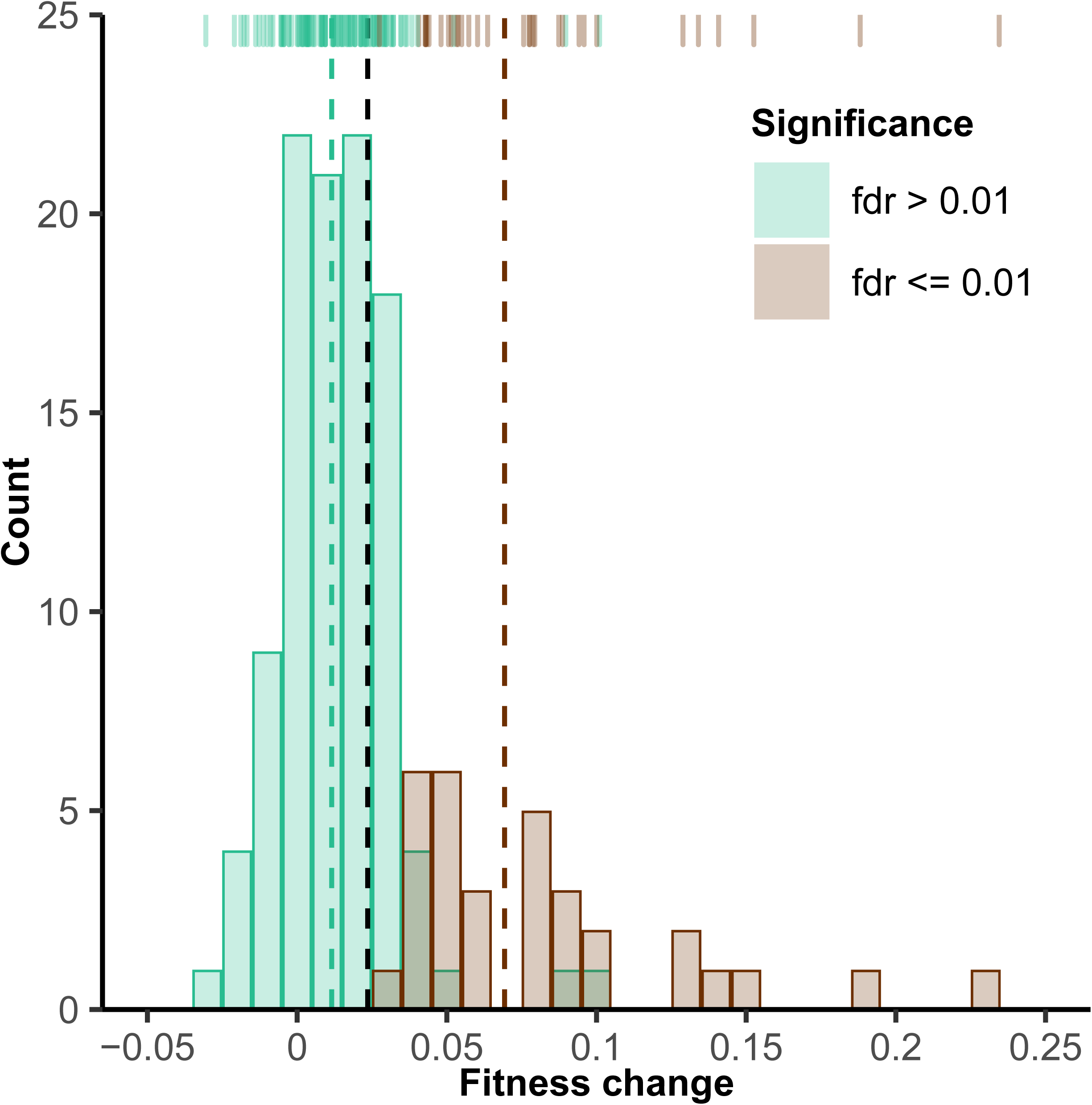
Change in fitness over 250 generations of experimental evolution. Histogram depicts fitness increase quantified by competition against a common reference strain. Fitness increase is generation-250 fitness minus generation-0 fitness. Orange and green bars indicate strains with fitness values significantly greater or not significantly greater than their own fitness at generation 0 at a false discovery rate of 1%, respectively. No strains significantly decreased in fitness. 16/152 entries with large standard error (> 0.029) among replicates are not displayed, but are included for all statistical testing. Dashed lines indicate the means of each group, with black indicating the overall mean.

Next, we evaluated the effect of evolutionary treatment on fitness change. We found significant variation in fitness among treatments (P < 10e-6, Figure 5, Supplemental Table S11). Relative to our standard culture conditions, diploids in CM (“no stress”) with 1:1000 transfer dilution, we found greater (more positive) fitness change in diploid strains evolved in salt stress (P < 10e-6), in haploid strains evolved in standard culture conditions (P = 0.01), and in diploid strains evolved with a less extreme (1:250) daily transfer dilution (P = 9.49e-4). Relative to the CM (“no stress”) diploid treatment (our standard culture conditions), we found less (less positive) fitness change in diploid strains evolved in ethanol stress (p = 3.99e-3). We found no significant effect of a more extreme (1:4000) daily transfer dilution on fitness change (p=0.39). In addition to the treatment effects, there was also a negative effect of barcode cross contamination rate on fitness change (P < 10e-6).

**Figure 5.**
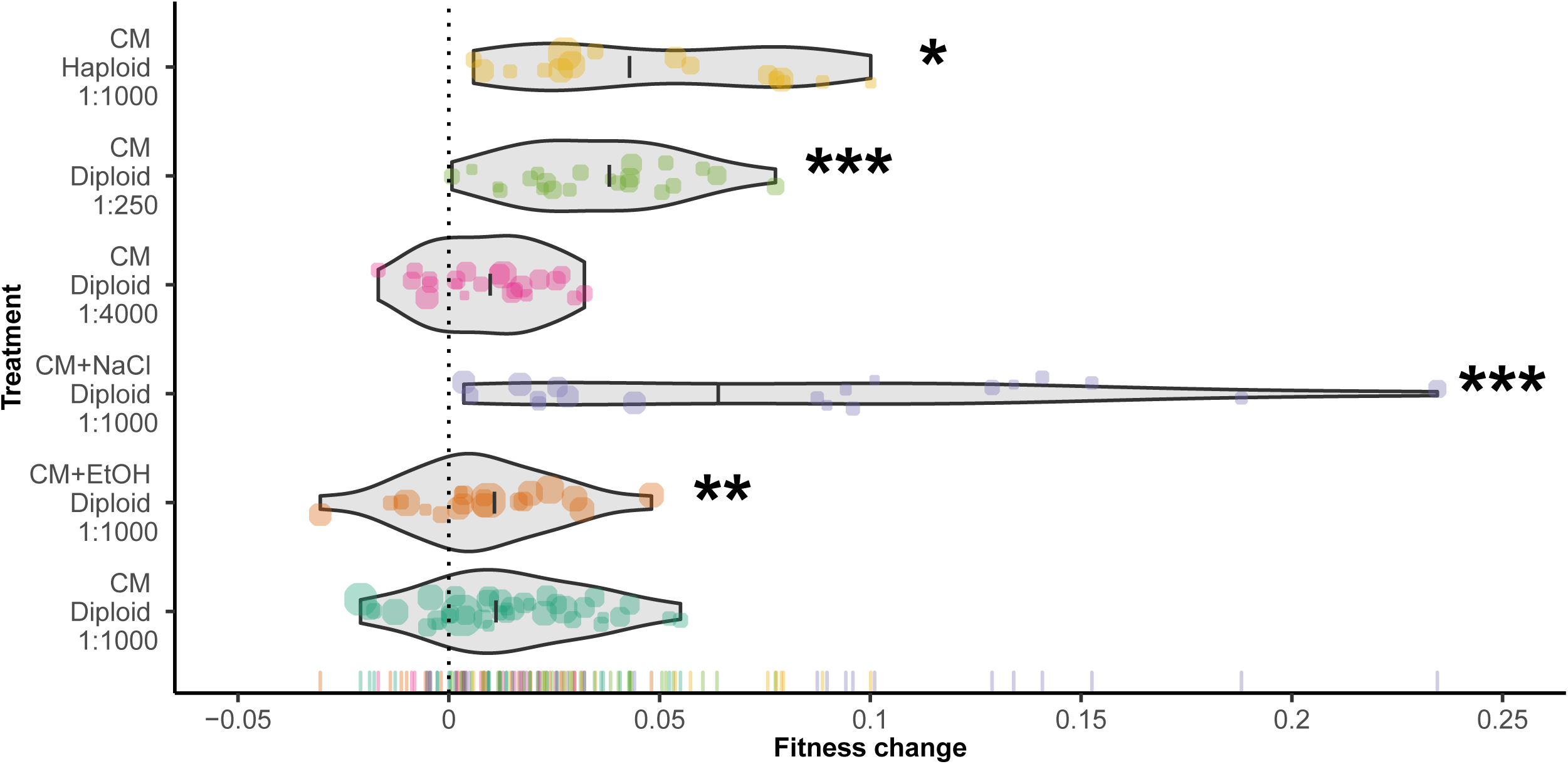
Change in fitness across six treatments in 250 generations of experimental evolution. Violin plots show the density of 152 yeast strains’ change in fitness with treatment labels indicating the medium, ploidy and dilution rate. Points indicate mean fitness change across 4 replicates for individual barcodes; point sizes reflect the number of reads underlying each datapoint and colors indicate evolutionary treatments. Treatment mean fitness changes are depicted as heavy black crossbars. 16/152 entries with large standard error (> 0.029) among replicates are not displayed, but are included for all statistical testing. Treatments significantly different from the control treatment, diploid yeast evolved under a standard 1:1000 transfer dilution in CM, in the associated mixed-effects model are marked with asterisks.

#### Evolutionary dynamics

In experimental evolution, adaptation can influence the relative abundance of barcodes evolving in competition (Kao and Sherlock 2008; Selmecki et al. 2015). In addition to characterizing fitness change for the evolved barcodes, we tracked the relative proportions of the barcoded-yeast pairs at seven timepoints during our evolution experiment to examine the evolutionary dynamics generated by adaptation and other processes (e.g., drift). We collected 8 measurements from these evolutionary dynamics: (1) barcode fixation, (2) the time-point and (3) magnitude of the maximum change in relative abundance in comparison to the starting conditions, (4) the time-point and (5) magnitude of the maximum rate of change between adjacent time-points, (6) the time-point and (7) magnitude of the maximum difference in BC proportions, and (8) the total cumulative change in barcoded relative abundance summed across all time-points.

Out of the seven measures of adaptive dynamics that were amenable to statistical testing (i.e., all adaptive dynamics measures other than fixation rate, which had sparse imbalanced binary data), six significantly differed by treatment (Table 1; see Supplemental Figures S3:S9 and Supplemental Tables S12:S18 for individual adaptive dynamics plots and results tables, respectively). Relative to the CM-diploid treatment (our standard culture conditions) both the salt and haploid treatments were associated with a larger maximum change in relative abundance in comparison to the start (p = 5.24e-5, p = 5.85e-3, respectively) and a more extreme maximum difference in barcode proportion (p = 2.77e-4, p = 6.01e-6, respectively). The salt treatment was also associated with an earlier time-point of the maximum change in abundance relative to the start (p = 0.01), and the haploid treatment was associated with an earlier maximum rate of change (p = 1.35e-5). The total cumulative change in barcode abundance was greater for the 1:4000 dilution treatment than the 1:1000 dilution treatment (p = 1.77e-3). Total cumulative change in barcode abundance was significantly less for haploids than diploids (p = 4.9e-2), and significantly less in the ethanol stress treatment than in the CM with no stress treatment (p = 2.98e-2). Finally, we observed barcodes that approached fixation in eleven percent (9/76) of our populations. Most fixation events were observed in the salt (3/11; 27.3%) and haploid (5/11; 45.5%) treatments, one instance was found in the CM-diploid treatment and no instances of fixation were observed in the ethanol, 1:4000 dilution, nor 1:250 dilution treatments.

**Table 1.**
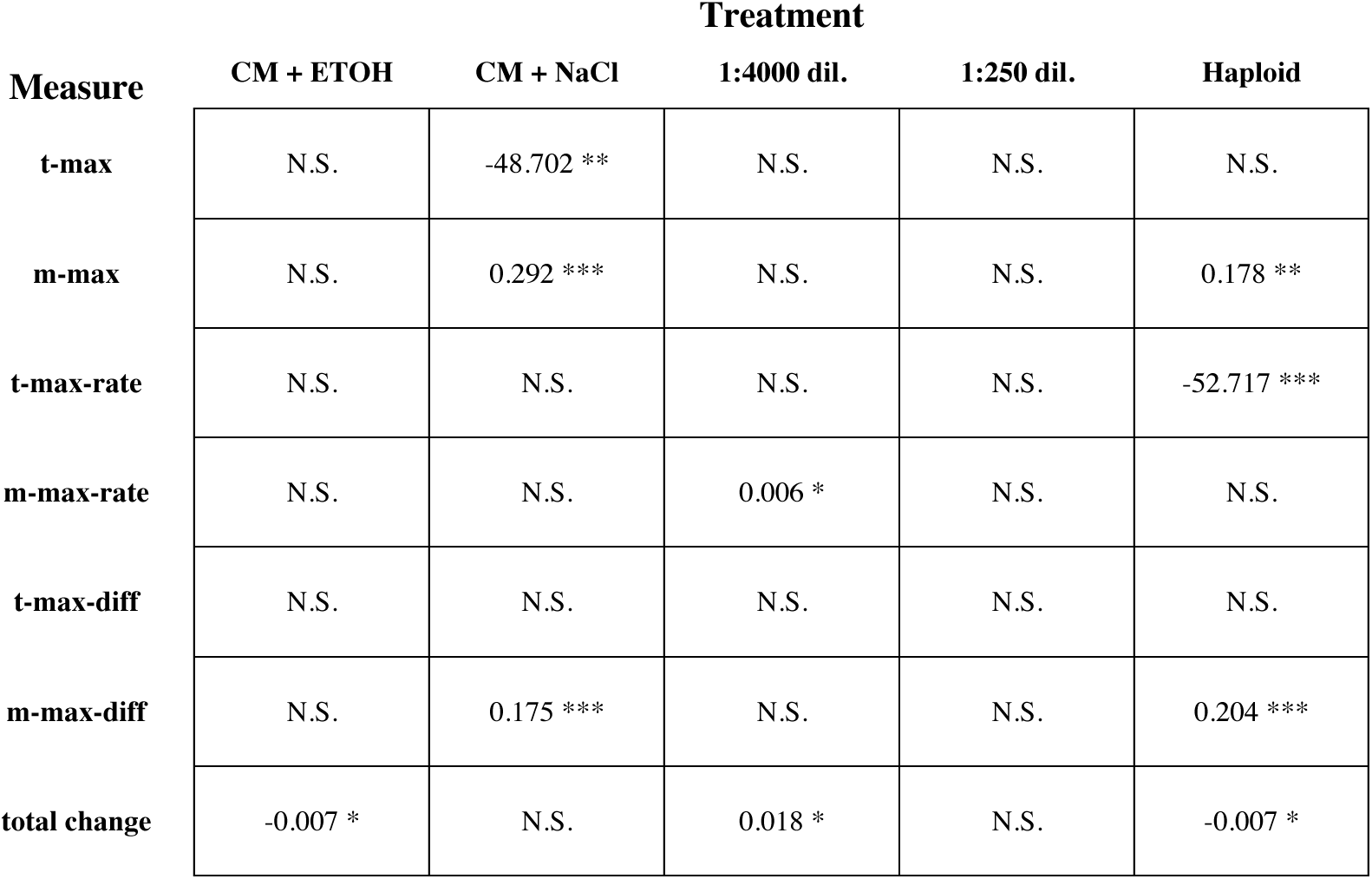
Associations between evolutionary dynamics and evolutionary treatments. Estimates for significant differences between each evolutionary treatment and the control (CM-diploid, 1:1000 transfer) are shown for seven different measures of barcoded dynamics (t-max, m-max, t-max-rate, m-max-rate, t-max-diff, m-max-diff, and total change).

| Measure | Treatment |  |  |  |  |
| --- | --- | --- | --- | --- | --- |
|  | CM + ETOH | CM + NaCl | 1:4000 dil. | 1:250 dil. | Haploid |
| t-max | N.S. | -48.702 ** | N.S. | N.S. | N.S. |
| m-max | N.S. | 0.292 *** | N.S. | N.S. | 0.178 ** |
| t-max-rate | N.S. | N.S. | N.S. | N.S. | -52.717 *** |
| m-max-rate | N.S. | N.S. | 0.006 * | N.S. | N.S. |
| t-max-diff | N.S. | N.S. | N.S. | N.S. | N.S. |
| m-max-diff | N.S. | 0.175 *** | N.S. | N.S. | 0.204 *** |
| total change | -0.007 * | N.S. | 0.018 * | N.S. | -0.007 * |

## Discussion

Experimental evolution has proven to be a valuable approach for studying a range of evolutionary questions. In this study we implemented a genetic barcoding system in *S. cerevisiae* to increase the efficiency and throughput of measuring fitness. Through barcoding we were able to employ a complex experimental design, increase throughput of fitness measurement by multiplexing, and provide a relatively simple means to track the evolutionary dynamics of barcodes evolving in competition during the experimental evolution period. Accordingly, we have demonstrated the potential for detecting fitness differences in six experimental evolution treatments and have shown that, while the typically low levels of barcode cross-contamination we observe cannot be completely eliminated, their effects on inference can be minimized through simple statistical procedures. Below we discuss the merits and drawbacks of the barcoding system, and its capabilities for efficient and high throughput quantification of fitness changes that occur during experimental evolution.

One important consideration for barcoding systems like the one we examine here and the system described by Levy, Blundell, and colleagues (Levy et al. 2015), is that barcoded strains are not necessarily identical to one another at the beginning of an experiment (i.e., have equal fitness) even when all barcoded variants are produced from a single ancestral clone. Although we found no significant fitness differences among the majority of barcoded strains, we note that we did indeed observe a few strains with significant deviations from the population mean fitness. Given the location of our barcode insertions (i.e., a non-functional region of the genome) it is unlikely that the barcodes themselves generated these fitness differences. A perhaps more likely explanation is that these differences arose from mutations that occurred during transformation (Giaever and Nislow 2014) or shortly thereafter. Regardless, this limitation could be mitigated by (1) removing strains with unequal generation-0 fitness entirely from their analyses, (2) quantifying initial fitness differentials and including them as a covariate in downstream analyses, or (3) looking explicitly at change in fitness for each individual barcode as we do here. We note that initial fitness differentials may be of interest themselves, given that they can potentially impact evolutionary outcomes (Barrick et al. 2010; Kryazhimskiy et al. 2014; Jerison et al. 2017) and the types of mutations that successfully spread through an experimental population (MacLean, Perron, and Gardner 2010).

In addition to throughput, the power to detect small changes in fitness in evolved strains is a critical parameter of any experimental evolution system. We estimated 80% power to detect a fitness difference of 1.2% in the proof-of-concept assays and 80% power to detect a fitness change of 3.0% in the 250-generation evolution experiment for any individual barcode. While a formal power analysis is not often reported, a review of the literature suggests that prior studies have detected fitness differences between 0.5% and 5% (Gresham et al. 2008; Lang, Botstein, and Desai 2011; McDonald, Rice, and Desai 2016; Venkataram et al. 2016; Fisher et al. 2018; Marad, Buskirk, and Lang 2018). For example, one previous barcoded fitness assay found between replicate deviations of 1-2% and between batch deviations of up to 5% (Venkataram et al. 2016). Some studies report confidence intervals significantly lower than 0.5%, but these studies tend to employ a very large number of replicates (MacLean, Perron, and Gardner 2010; Gallet et al. 2012; Kryazhimskiy, Rice, and Desai 2012). We also note that our power to detect treatment effects was much higher than our ability to detect changes in fitness for individual barcoded strains. Specifically, we estimated 99.8% power to detect fitness differences of 1% between treatments using the error in our proof-of-concept fitness assays, and 87.7% power to detect fitness differences of 1% between treatments using the actual 250-generation experiment data, indicating that the power to detect treatment effects in this system is similar to traditional systems that utilize competition-based assays to quantify fitness (Lang, Botstein, and Desai 2011; McDonald, Rice, and Desai 2016; Fisher et al. 2018; Marad, Buskirk, and Lang 2018). We note that power to detect small fitness effects can be increased by measuring the log-linear slope of barcode frequency change across several time-points and by accounting for PCR duplicates when estimating barcode frequencies (Blundell and Levy 2014; Venkataram et al. 2016; F. Li, Salit, and Levy 2018). While these strategies were not employed here, they could be incorporated into our system.

While the error measured in our fitness assays is consistent with most prior studies, pooled fitness assays do have additional sources of error. Prior studies using pooled fitness assays estimated that pools containing 1000 cells per strain are expected to have an error rate of 4.4% (Pierce et al. 2007) after two rounds of dilution due to sampling error. The expected number of cells per barcoded strain at inoculation in our system is 9,000 (3.0×10^8 cells/ml at saturation in 0.6ml of media, diluted 1:1000 and divided by up to 20 strains per assay). However, the number of cells could be much lower due to sampling or to low frequency of the barcode of interest in a well at the end of the 250 generations of evolution. Lineages with large changes in fitness are expected to deviate the most from equal numbers of cells in a well. Consistent with this possibility, the RMSE of fitness in generation 250 (4.34) is higher than the RMSE of fitness in generation 0 (2.92), where barcode frequencies are much more uniform. This provides one explanation for our finding that error among replicates is associated with the magnitude of the fitness change. Thus, while there are some limitations to pooled fitness assays, these limitations can be compensated by examination a modest number of strains per treatment –as done in this study– or by increasing the number of replicate assays for each individual strain.

Another important consideration when designing an experimental evolution system is that it must be robust to contamination. While culture contamination is rare in experimental evolution (Lenski et al. 1991; T. F. Cooper and Lenski 2010), barcode cross-contamination is possible. Our results are consistent with prior work on this topic. Specifically, while we found no evidence of culture contamination in any growth blanks, we detected a uniformly low level of diffuse barcode cross-contamination in multiplexed fitness assays and evolutionary dynamics samples. Because there was no conspicuous pattern of cross-contamination, we suggest that the observed contamination was likely introduced during sample preparation and/or DNA sequencing. DNA extraction is a likely source of cross-contamination in samples processed in strip-tubes or 96-well plate formats that prioritize throughput. However, we minimized the chance for contamination in the DNA isolation step in our experiments by isolating DNA with individual reaction tubes for each sample. Furthermore, contamination in the DNA extraction step would not be consistent with the low-level diffuse contamination observed, a pattern that tends to be more indicative of contamination occurring downstream of DNA extraction. Several other sources of contamination are possible, including primer contamination during the index addition for PCR (Lo, Mehal, and Fleming 1988) and index switching during library construction or sequencing (Illumina 2017; Sinha et al. 2017; Costello et al. 2018). The latter possibility seems nevertheless less likely because all PCR steps were performed separately prior to pooling of libraries and because no previous studies report index switching for IonTorrent sequence data to our knowledge.

The strength and efficiency of barcoded experimental evolution is further evidenced by the biological results of this study. Here, barcoding enabled us to evolve a moderate number of strains (152) for 250 generations across six treatments, and to conduct a total of 532 evolutionary dynamics measurements and 1,216 fitness assays related to these manipulations in a relatively short amount of time and with limited resources.

Reassuringly, our biological results are largely consistent with prior work. We find greater fitness increases, i.e., a greater rate of adaptation, in haploids than diploids. This finding agrees with another study that found faster rates of adaptation and larger effective population sizes in haploids relative to diploids (A. C. Gerstein et al. 2011). In a related study, Selmecki et al., (Selmecki et al. 2015) found faster adaptation in tetraploids than diploids, but no difference in rate of adaptation between haploids and diploids, potentially suggesting a trend opposite to ours of increasing adaptive rate with increasing ploidy. Further exploration of the evolutionary differences between haploid and diploid yeast reveals more support for the observation that diploids tend to evolve slower than haploids (Fisher et al. 2018; Marad, Buskirk, and Lang 2018), and indicates that haploids and diploids have different targets and types of mutations as well as access to differing spectrums of beneficial mutations (Fisher et al. 2018; Marad, Buskirk, and Lang 2018). However, differences between haploids and diploids must be treated with caution because there is mounting evidence that haploid yeast evolves towards diploidy under experimental evolution conditions (Aleeza C. Gerstein and Otto 2011; R. Dhar et al. 2011; Selmecki et al. 2015), potentially due to an initial spike in fitness associated with autodiploidy (Fisher et al. 2018). We also note that neither haploids or diploids were put through a sexual cycle; allowing for sexual reproduction speeds the rate of adaptation (McDonald, Rice, and Desai 2016). We also find greater fitness increase in complete medium (CM) plus NaCl stress than in CM alone, which was not surprising given what is known about adaptation to NaCl stress in *S. cerevisiae* (Blomberg 1995; R. Dhar et al. 2011; Park, Yang, and Kim 2015; Tekarslan-Sahin, Alkim, and Sezgin 2018). In contrast, we were surprised to detect less fitness increase in CM plus EtOH stress than in CM alone. There are several non-mutually exclusive explanations for this result. It is possible that ethanol did not present a significant stress (selective pressure) to the cells once they had attained physiological adaptation to the medium, i.e., acclimation (Huang et al. 2018). It is also possible that adaptations to CM and adaptations to ethanol exhibit antagonistic pleiotropy, similar to what has been found in experiments contrasting rich and poor media (Minty et al. 2011) or exploring adaptation to other chemical stressors (Reyes, Abdelaal, and Kao 2013). Pleiotropy could also shed light on the marked adaptation observed in the NaCl treatment given that adaptation to CM and NaCl stress may exhibit complementarity via synergistic or positive pleiotropy (Ostman, Hintze, and Adami 2012; Riddhiman Dhar et al. 2013; McGee et al. 2016; K. A. Hughes and Leips 2017). We find no difference in fitness change between our standard (1:1000) dilution treatment and a treatment with a more extreme (1:4000) daily transfer dilution. We do, however, find a greater increase in fitness when a less extreme (1:250) daily transfer dilution was used instead of the standard (1:1000) dilution. While these transfer dilution findings are not necessarily expected (Gerrish and Lenski 1998; De Visser et al. 1999; De Visser and Rozen 2006; Kryazhimskiy, Rice, and Desai 2012), they nevertheless support earlier findings that less extreme bottlenecks favor the maintenance of adaptive mutants (Wahl, Gerrish, and Saika-Voivod 2002) and that large populations are less adaptively constrained than small ones in simple environments (De Visser and Rozen 2006). They are also consistent with previous evidence for greater mean fitness increase in wide (relatively unrestricted) versus narrow (relatively restricted) bottleneck populations (Schoustra et al. 2009).

In addition to high-throughput fitness assays, barcoding enabled us to track the relative proportions of barcoded lineage pairs evolving in competition during the experimental evolution period (Hegreness et al. 2006; Blundell and Levy 2014; Levy et al. 2015; V. S. Cooper 2018). As expected, we found general agreement between the evolutionary dynamics results and endpoint fitness assay results in our six experimental treatments. Our haploid and NaCl stress treatments both displayed dynamics consistent with more extreme increases in fitness, including greater change in barcode abundance relative to the starting conditions and a greater maximum difference in barcode abundance than diploids in CM. Interestingly, haploids also showed signs of earlier adaptation than diploid strains in similar conditions, as evidenced by an earlier generation of maximum rate of change in barcode abundance (also noted in (Blundell and Levy 2014; Levy et al. 2015; Selmecki et al. 2015)) and a lower total change in barcode abundance over 250 generations. Although this latter result may seem paradoxical, it is consistent with the observation that haploids adapt earlier than diploid strains (A. C. Gerstein et al. 2011), and is expected if a greater proportion of the total change in barcode abundance of haploids in our experiments happened in the first 100 generations (before we collected any evolutionary dynamics data).

Despite significant differences in barcode dynamics, there are some limitations to interpreting these results. Because we mostly assessed abundance every 50 generations, it is possible that we missed some of the adaptive dynamics; dense temporal sampling is ideal for a full picture of evolutionary dynamics in populations subjected to experimental evolution (Hegreness et al. 2006). Furthermore, due to projected read depth constraints, barcode frequencies over time were not measured in replicate for the 250-generation evolution project. Finally, changes in barcode abundance over long time scales may not be directly related to final fitness as these measures are calculated over 250 generations of evolution, during which relative fitness relationships between evolving barcodes could change significantly. For example, the 1:4000 transfer dilution bottleneck treatment had elevated rates of change in barcode abundance and a high amount of total change in barcode abundance without a concomitant increase in fitness, possibly as a result of the potentially strong effects of drift when severe bottlenecks are frequent. We suggest that future studies employ a denser and more even longitudinal-evolutionary dynamics sampling scheme, with replication, to maximize the value and precision of this type of lineage tracking or evolutionary dynamics data.

## Conclusions

In summary, we conclude that the barcoded yeast system that we describe here offers a flexible, yet high-throughput means of fitness measurements and provides a relatively simple means for lineage tracking. These two characteristics support our ability to design more complex and potentially more informative evolutionary experiments. We note that although barcode cross-contamination imposes some limitations on the implementation of this system, it is possible to track the origin and rates of such contamination and, therefore, to statistically consider its effects on experimental outcomes. While prior barcoding systems leverage massive lineage tracking over short time-periods (Blundell and Levy 2014; Levy et al. 2015; Venkataram et al. 2016; Y. Li et al. 2018), our system uses far fewer barcodes but is more flexible at handling different experimental designs, such as the multiple treatments that we employed in this study. Thus, we conclude that this system represents an informative step towards the design and implementation of experimental evolution systems capable of both long-term evolution studies and high throughput fitness measurements.

## Supporting information

Supplement Descriptions

## Acknowledgements

The authors thank the members of the Carlos A. Botero and Justin C. Fay labs, specifically Angela Chira, Trevor Fristoe, Emery Longan, and Dilys Vela for ongoing feedback and helpful advice.

