## Supplement Descriptions for "High-throughput analysis of adaptation using barcoded strains of *Saccharomyces cerevisiae*"

**Supplemental File Descriptions.**

**File Name: Supplemental Table S1**

File Format: Word Document, .docx

Title of Data: Strain Construction, Yeast strains and inserted MOBY barcode sequences.

Description of Data: Strain construction -- Oligo Name indicates the name of the forward or reverse primer; Oligo Names that contain numbers preceded by an "R" are reverse primers. Full Sequence notes the full Oligo sequence in the 5'-3' direction (top). MOBY barcodes -- Columns 1 and 2, resp., contain diploid and haploid yeast strain ID's. Only yeast strains used in the current project are included (diploids, n = 92; haploids, n = 13). Columns 3 and 4, resp., contain the MOBY barcode uptag (5'-3') and MOBY barcode downtag (3'-5-) sequences that uniquely identify the haploid and diploid yeast strains in the same row (bottom).

**File Name: Supplemental Table S2**

File Format: Word Document, .docx

Title of Data: Treatments for 250-generation experimental evolutions.

Description of Data: Treatment ID is a numerical ID for each of six treatments; Evo. Wells is the number of populations assigned to each treatment; Barcodes per Evo. Well is the number of sympatric barcodes per evolutionary population; Ploidy, Evo. Medium, and Evo. Transfer Dilution indicate the yeast strain ploidy, evolutionary medium type, and daily passage dilution for each treatment, respectively.

**File Name: Supplemental Table S3**

File Format: Word Document, .docx

Title of Data: Ion Proton sequencing primers.

Description of Data: Oligo Name indicates the name of the forward or reverse primer; Oligo Names that contain numbers preceded by an "R" are reverse primers. Full Sequence notes the full Oligo sequence in the 5'-3' direction. Barcode sequence shows the unique genetic barcode within each primer, in the 5'-3' direction, used in multiplexing and demultiplexing libraries.

**File Name: Supplemental Table S4**

File Format: Excel document, .xlsx

Title of Data: Estimated fitness change in 250 generations evolution and significance of 91 barcoded yeast strains.

Description of Data: Summary of results for a fixed-0-intercept linear model of mean-corrected barcoded yeast initial fitness. Predictors are individual barcoded yeast strain ID’s. FDR adjusted p-values for a 2-tailed test are reported.

**File Name: Supplemental Table S5**

File Format: html document, .html

Title of Data: Data table of power calculations using error in the POC fitness assays data to support figure 3 panel A.

Description of Data: Data used to generate figure 3 panel A (green points). Columns, in order, are: fitness change (fitchange), population standard deviation in the POC fitness assays data among replicate measures of fitness (psd), number of replicates per timepoint (n), power, and significance level (sig.level). fitchange and n are varied to calculate power; psd and sig.level are kept constant.

**File Name: Supplemental Table S6**

File Format: html document, .html

Title of Data: Data table of power calculations using error in the POC fitness assays data to support figure 3 panel B.

Description of Data: Data used to generate figure 3 panel B (green points). Columns, in order, are: degrees of freedom in the numerator (u), degrees of freedom in the denominator (v), power, effect size (f2), and significance level (sig.level). v and f2 are varied to calculate power; u and sig.level are kept constant.

**File Name: Supplemental Figure S1**

File Format: pdf document, .pdf

Title of Data: barcode cross-contamination rates for sequenced libraries.

Description of Data: A. Histogram of mean barcode contamination rate (as a percentage of total count) for the cross-contamination diagnostic samples included in the Proof of Concept Fitness Assays sequenced library. One datapoint is reported for each unique forward-reverse index pair (sample) in the sequenced library. B. Histogram of mean barcode contamination rate (as a percentage of total count) for samples included in the 250-generation experimental evolution project. For resequenced samples, only the less contaminated sample is retained. One datapoint is reported for each unique forward-reverse index pair (sample) in this consensus library.

**File Name: Supplemental Figure S2**

File Format: pdf document, .pdf

Title of Data: Decreased contamination after reprocessing samples.

Description of Data: Violin plots showing decrease in barcode contamination rate for a set of samples from the 250-generation evolution experiment that were DNA extracted, PCR amplified, and sequenced two separate times. Colors depict sequencing runs: cyan for run 1., and orange for run 2. Point sizes reflect the number of reads underlying each datapoint. Mean contamination rate for run 1., and run 2., are depicted as heavy black crossbars.

**File Name: Supplemental Table S7**

File Format: html document, .html

Title of Data: Data table of power calculations using error in the change in fitness data for the 250-generation experiment to support figure 3 panel A.

Description of Data: Data used to generate figure 3 panel A (orange points). Columns, in order, are: fitness change (fitchange), population standard deviation in the POC fitness assays data among replicate measures of fitness (psd), number of replicates per timepoint (n), power, and significance level (sig.level). fitchange and n are varied to calculate power; psd and sig.level are kept constant.

**File Name: Supplemental Table S8**

File Format: html document, .html

Title of Data: Data table of power calculations using error in the change in fitness data for the 250-generation experiment to support figure 3 panel B.

Description of Data: Data used to generate figure 3 panel B (orange points). Columns, in order, are: degrees of freedom in the numerator (u), degrees of freedom in the denominator (v), power, effect size (f2), and significance level (sig.level). v and f2 are varied to calculate power; u and sig.level are kept constant.

**File Name: Supplemental Table S9**

File Format: html document, .html

Title of Data: Change in fitness over 250 generations of experimental evolution.

Description of Data: Summary of results for a fixed-0-intercept linear model of barcoded yeast fitness change over 250-generations of experimental evolution. Predictors are generation-250 barcoded yeast strains. FDR adjusted p-values for a 1-tailed (greater) test are reported.

**File Name: Supplemental Table S10**

File Format: Excel document, .xlsx

Title of Data: Standard error in change in fitness across six treatments in 250 generations of experimental evolution.

Description of data: summary of results for linear model of standard error in change in barcoded yeast strain fitness over 250-generations of experimental evolution.

**File Name: Supplemental Table S11**

File Format: html document, .html

Title of Data: Change in fitness across six treatments in 250 generations of experimental evolution.

Description of Data: summary of results for linear mixed effects model of change in barcoded yeast strain fitness over 250-generations of experimental evolution.

**File Name: Supplemental Table S12**

File Format: html document, .html

Title of Data: t-max by treatment model results.

Description of Data: summary of results for linear mixed effects model of generation of maximum deviation in barcode abundance from initial conditions over 250-generations of experimental evolution by treatment.

**File Name: Supplemental Figure S3**

File Format: pdf document, .pdf

Title of Data: t-max by treatment visualization.

Description of Data: Violin plot of generation of maximum deviation in barcode abundance from initial conditions for 152 yeast strains evolved across six evolutionary treatments for 250 generations. Point sizes reflect the number of reads underlying each datapoint and colors indicate evolutionary treatments. Treatment means are depicted as heavy black crossbars. Treatments significantly different from the control treatment are marked with an asterisk. The treatment with diploid yeast evolved under a standard 1:1000 transfer dilution in CM is selected as the reference level in this model.

**File Name: Supplemental Table S13**

File Format: html document, .html

Title of Data: m-max by treatment model results.

Description of Data: summary of results for linear mixed effects model of maximum deviation in barcode abundance from initial conditions over 250-generations of experimental evolution by treatment.

**File Name: Supplemental Figure S4**

File Format: pdf document, .pdf

Title of Data: m-max by treatment visualization.

Description of Data: Violin plot of magnitude of maximum deviation in barcode abundance from initial conditions for 152 yeast strains evolved across six evolutionary treatments for 250 generations. Point sizes reflect the number of reads underlying each datapoint and colors indicate evolutionary treatments. Treatment means are depicted as heavy black crossbars. Treatments significantly different from the control treatment are marked with an asterisk. The treatment with diploid yeast evolved under a standard 1:1000 transfer dilution in CM is selected as the reference level in this model.

**File Name: Supplemental Table S14**

File Format: html document, .html

Title of Data: t-max-rate by treatment model results.

Description of Data: summary of results for linear mixed effects model of generation of maximum rate of change in barcode abundance over 250-generations of experimental evolution by treatment.

**File Name: Supplemental Figure S5**

File Format: pdf document, .pdf

Title of Data: t-max-rate by treatment visualization.

Description of Data: Violin plot of generation of maximum rate of change in barcode abundance for 152 yeast strains evolved across six evolutionary treatments for 250 generations. Point sizes reflect the number of reads underlying each datapoint and colors indicate evolutionary treatments. Treatment means are depicted as heavy black crossbars. Treatments significantly different from the control treatment are marked with an asterisk. The treatment with diploid yeast evolved under a standard 1:1000 transfer dilution in CM is selected as the reference level in this model.

**File Name: Supplemental Table S15**

File Format: html document, .html

Title of Data: m-max-rate by treatment model results.

Description of Data: Summary of results for linear mixed effects model of magnitude of maximum rate of change in barcode abundance over 250-generations of experimental evolution by treatment.

**File Name: Supplemental Figure S6**

File Format: pdf document, .pdf

Title of Data: m-max-rate by treatment visualization.

Description of Data: Violin plot of magnitude of maximum rate of change in barcode abundance for 152 yeast strains evolved across six evolutionary treatments for 250 generations. Point sizes reflect the number of reads underlying each datapoint and colors indicate evolutionary treatments. Treatment means are depicted as heavy black crossbars. Treatments significantly different from the control treatment are marked with an asterisk. The treatment with diploid yeast evolved under a standard 1:1000 transfer dilution in CM is selected as the reference level in this model.

**File Name: Supplemental Table S16**

File Format: html document, .html

Title of Data: t-max-diff by treatment model results.

Description of Data: Summary of results for linear mixed effects model of generation of maximum difference in sympatric barcode abundance over 250-generations of experimental evolution by treatment.

**File Name: Supplemental Figure S7**

File Format: pdf document, .pdf

Title of Data: t-max-diff by treatment visualization.

Description of Data: Violin plot of generation of maximum difference in sympatric barcode abundance for 152 yeast strains evolved across six evolutionary treatments for 250 generations. Point sizes reflect the number of reads underlying each datapoint and colors indicate evolutionary treatments. Treatment means are depicted as heavy black crossbars. Treatments significantly different from the control treatment are marked with an asterisk. The treatment with diploid yeast evolved under a standard 1:1000 transfer dilution in CM is selected as the reference level in this model.

**File Name: Supplemental Table S17**

File Format: html document, .html

Title of Data: m-max-diff by treatment model results.

Description of Data: Summary of results for linear mixed effects model of magnitude of maximum difference in sympatric barcode abundance over 250-generations of experimental evolution by treatment.

**File Name: Supplemental Figure S8**

File Format: pdf document, .pdf

Title of Data: m-max-diff by treatment visualization.

Description of Data: Violin plot of magnitude of maximum difference in sympatric barcode abundance for 152 yeast strains evolved across six evolutionary treatments for 250 generations. Point sizes reflect the number of reads underlying each datapoint and colors indicate evolutionary treatments. Treatment means are depicted as heavy black crossbars. Treatments significantly different from the control treatment are marked with an asterisk. The treatment with diploid yeast evolved under a standard 1:1000 transfer dilution in CM is selected as the reference level in this model.

**File Name: Supplemental Table S18**

File Format: html document, .html

Title of Data: Total change in barcode abundance by treatment model results.

Description of Data: summary of results for linear mixed effects model of total (cumulative) change in sympatric barcode abundance over 250-generations of experimental evolution by treatment.

**File Name: Supplemental Figure S9**

File Format: pdf document, .pdf

Title of Data: Total change in barcode abundance by treatment visualization.

Description of Data: Violin plot of total (cumulative) change in sympatric barcode abundance for 152 yeast strains evolved across six evolutionary treatments for 250 generations. Point sizes reflect the number of reads underlying each datapoint and colors indicate evolutionary treatments. Treatment means are depicted as heavy black crossbars. Treatments significantly different from the control treatment are marked with an asterisk. The treatment with diploid yeast evolved under a standard 1:1000 transfer dilution in CM is selected as the reference level in this model.
