## Supplementary figures and images for "High-throughput analysis of adaptation using barcoded strains of *Saccharomyces cerevisiae*"

### Supplmental Figure 1

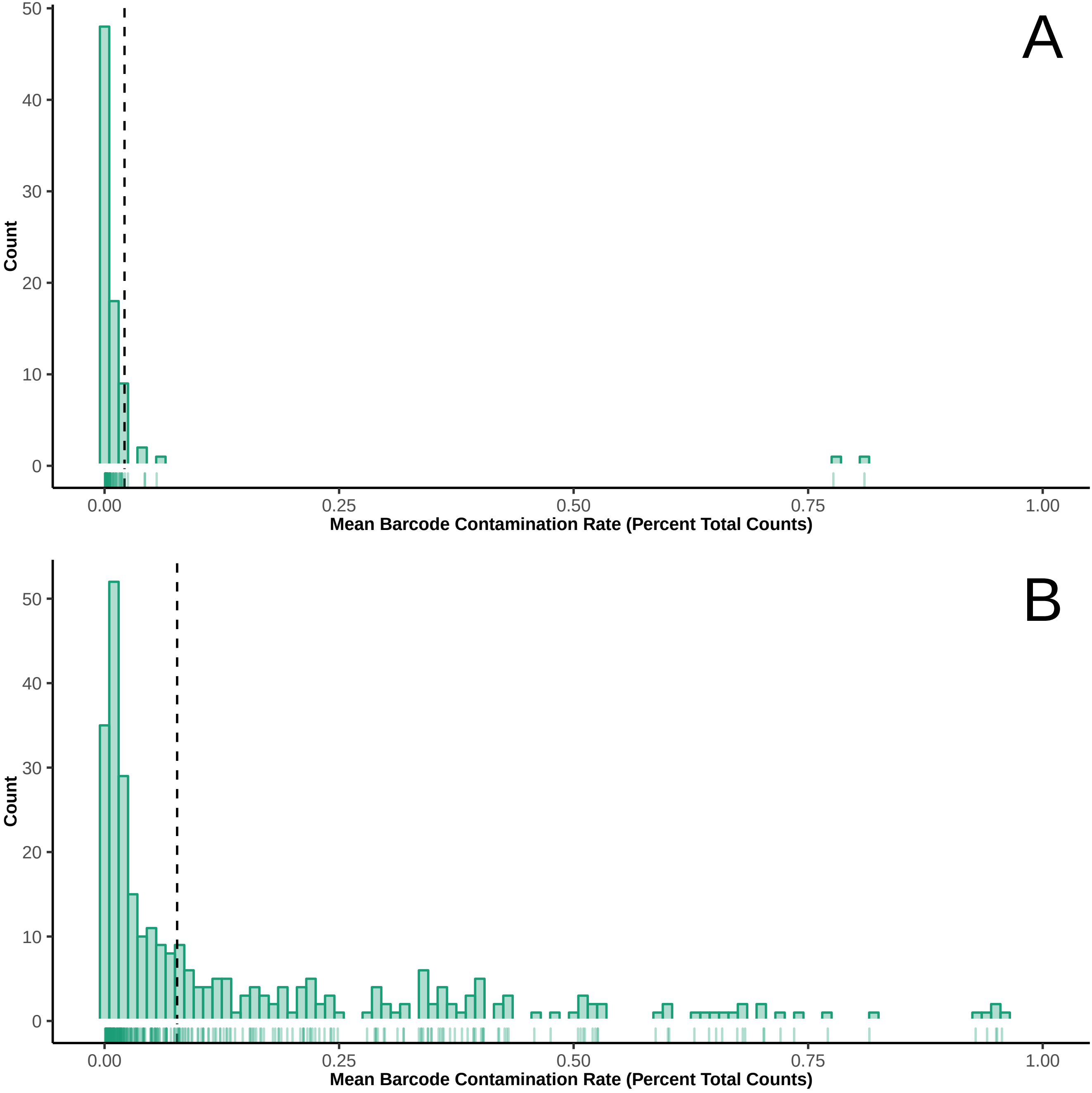

### Supplmental Figure 2

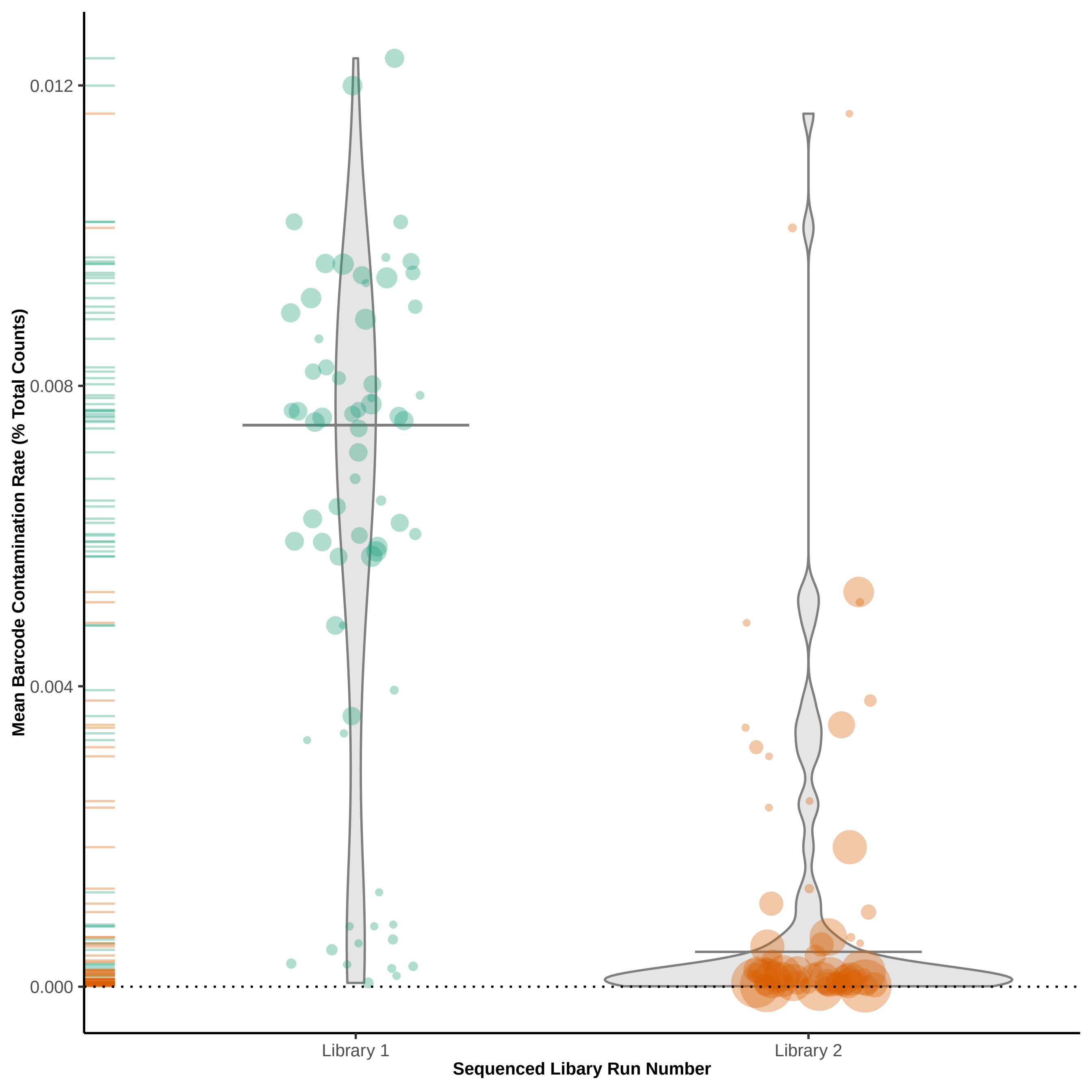

### Supplmental Figure 3

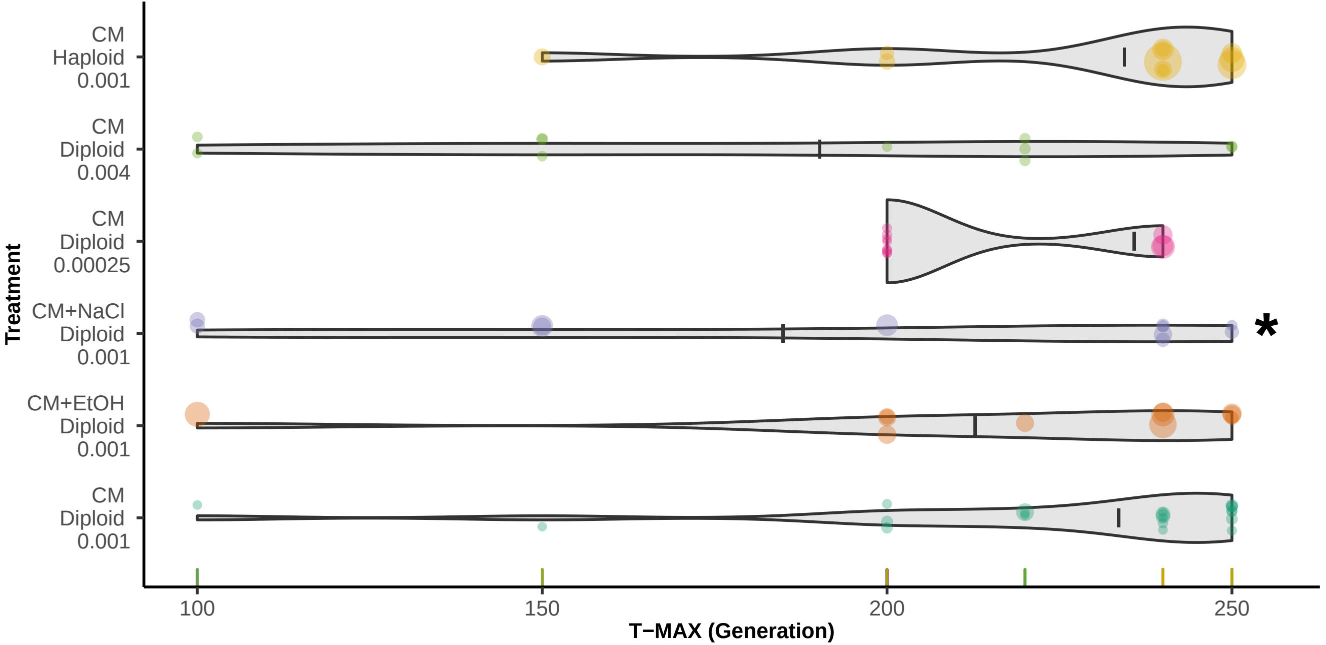

### Supplmental Figure 4

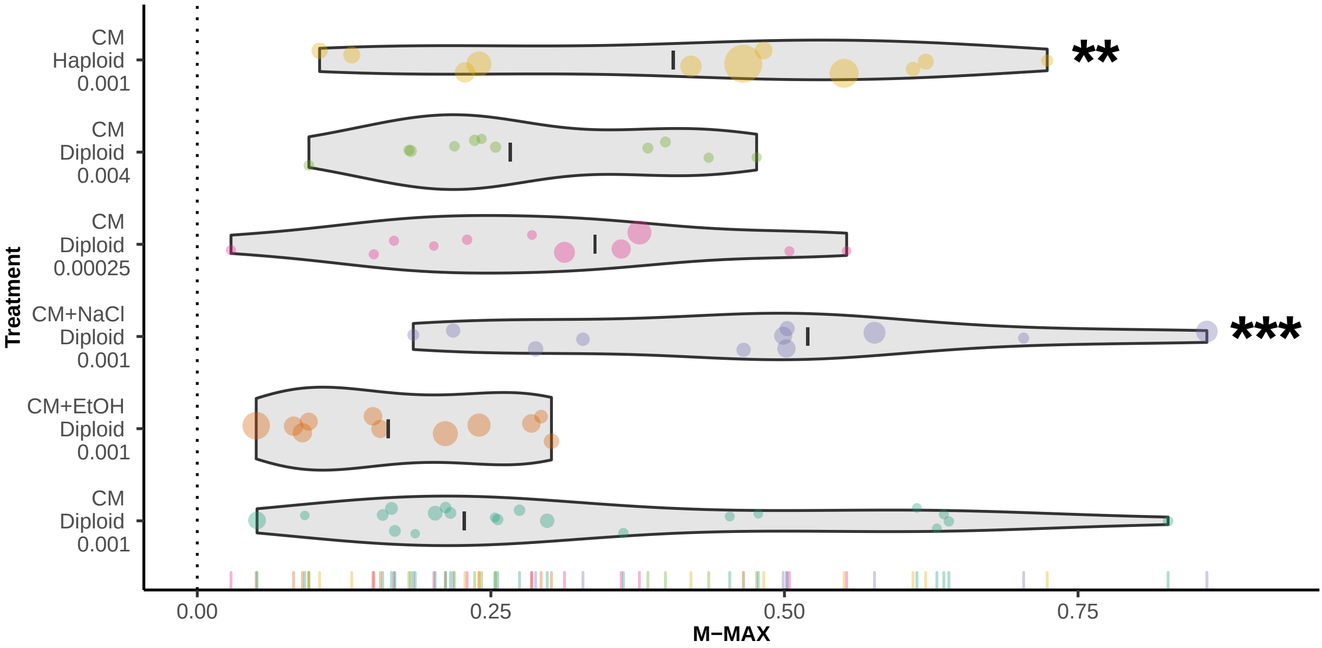

### Supplmental Figure 5

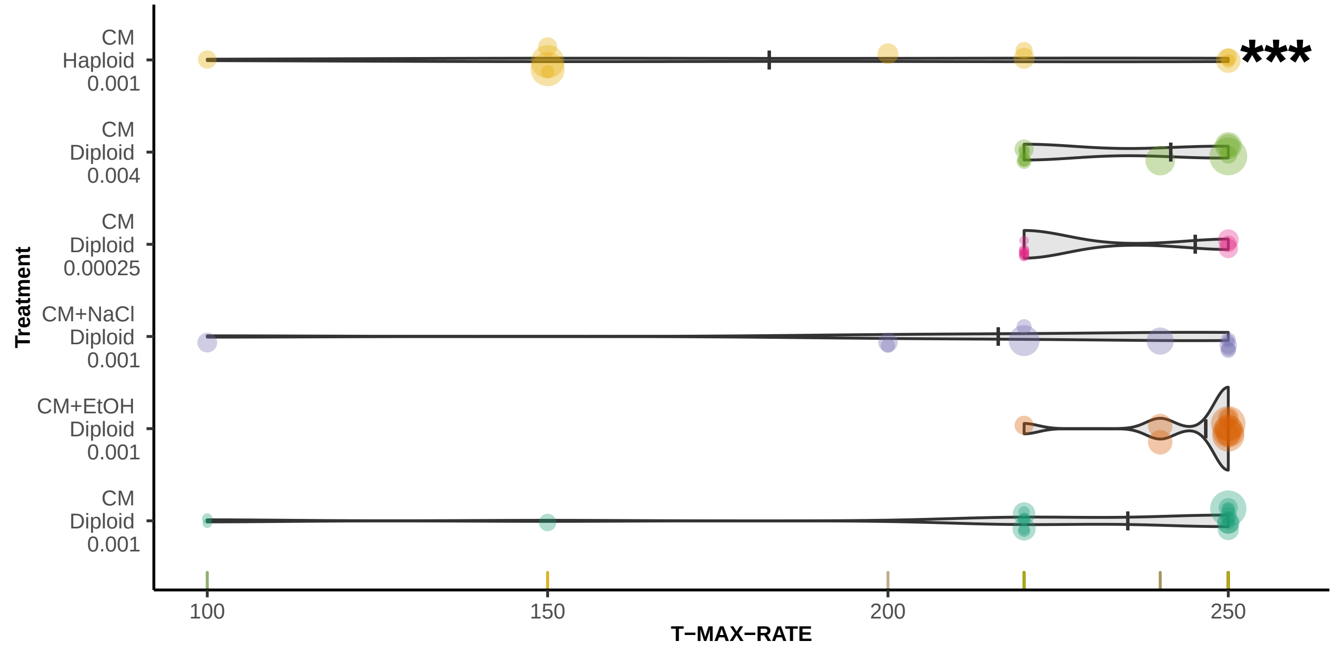

### Supplmental Figure 6

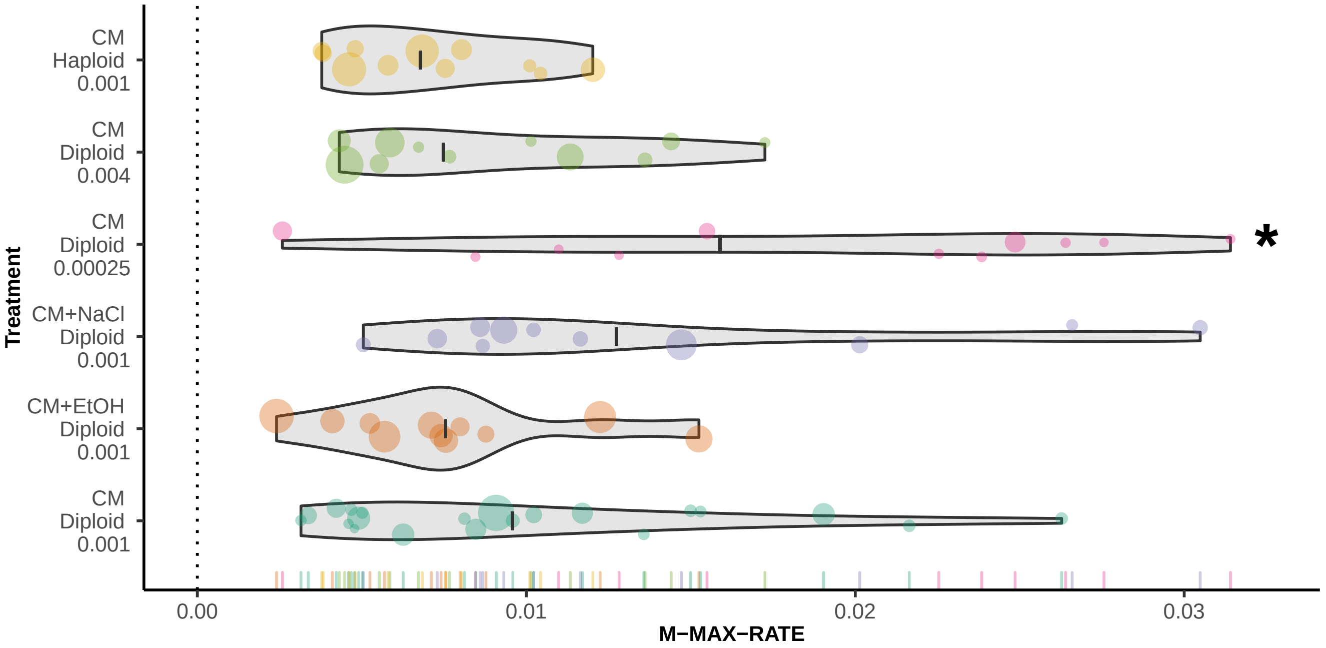

### Supplmental Figure 7

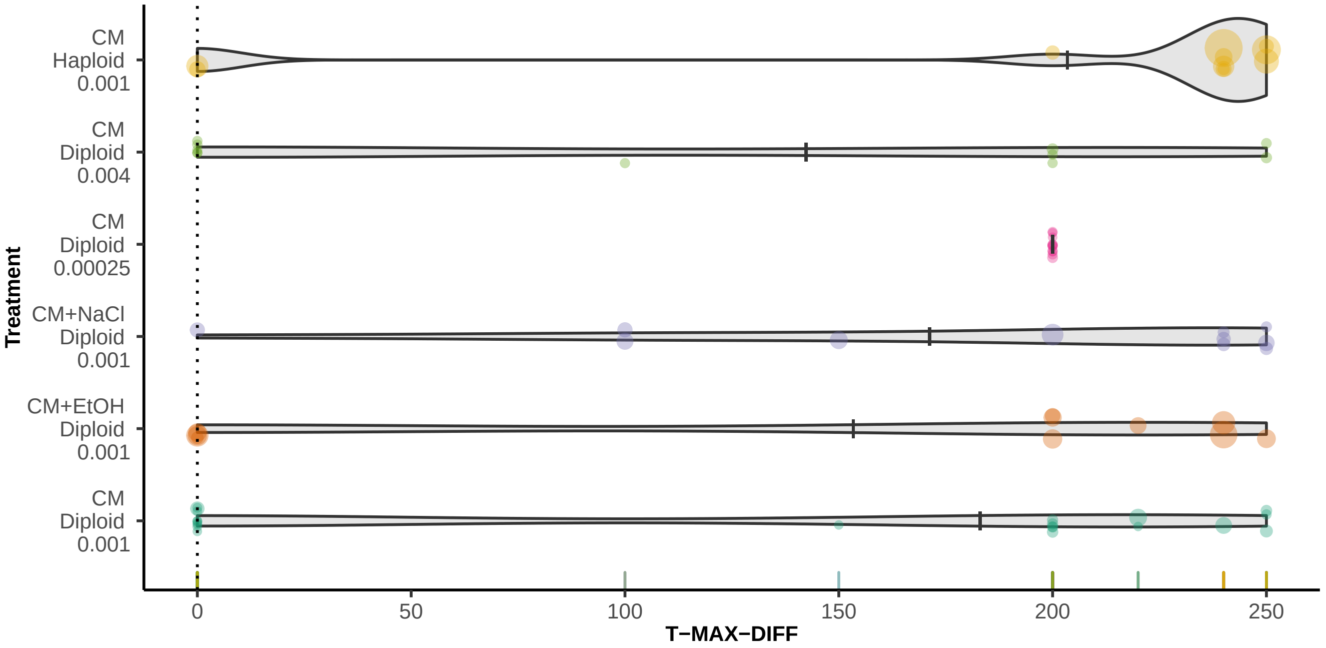

### Supplmental Figure 8

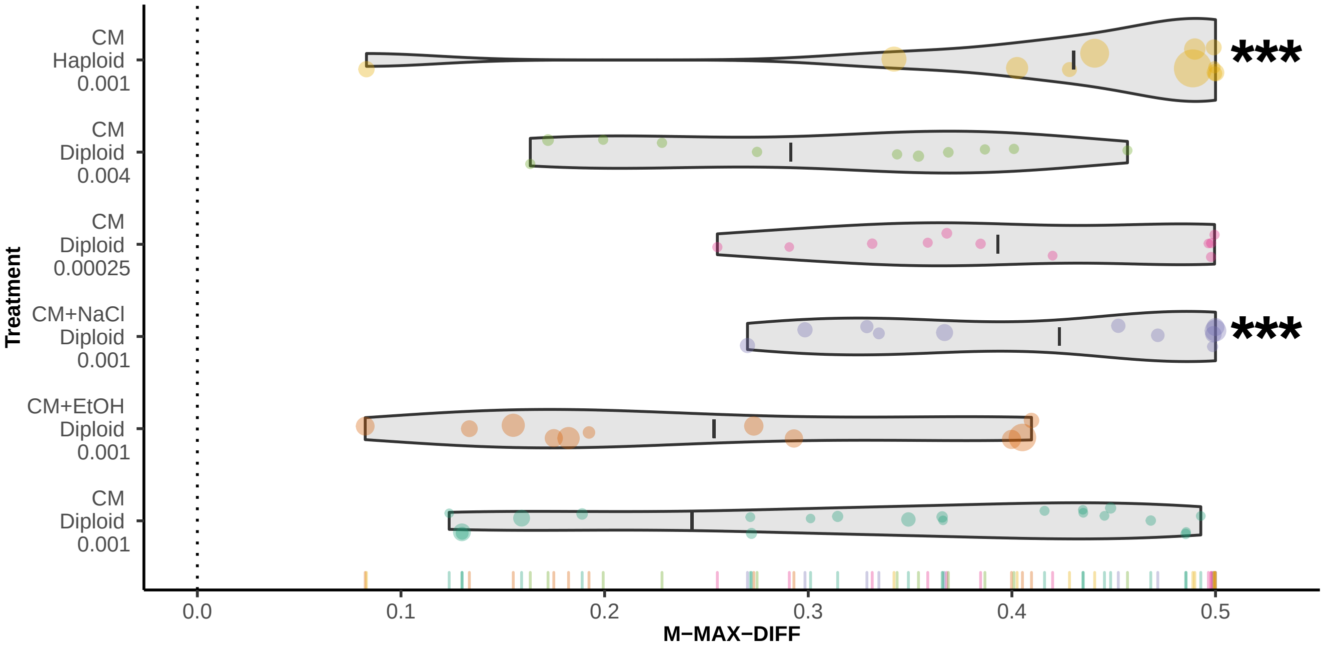

### Supplmental Figure 9

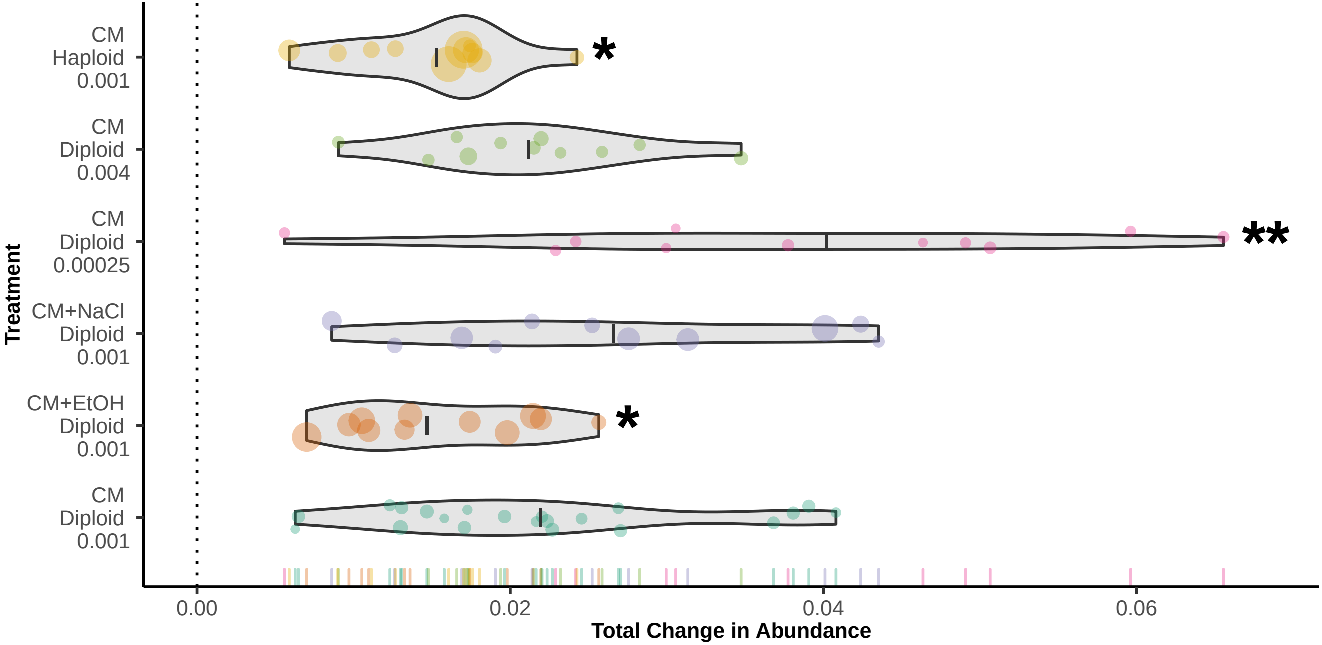
