## Supplementary material for "High-throughput analysis of adaptation using barcoded strains of *Saccharomyces cerevisiae*": Supplmental Table 1

| ***Oligo name*** | ***Full sequence (5'-3')*** |
| --- | --- |
| F2 | TCACAACAAATGGTCAACCATCAAGCACGGTAAAGCCAAGGATGTCCACGAGGTCTCT |
| R2 | GCGCAATGTCTTCGATGCCTTGAATCTCCAGTACGCTTTCCGGTGTCGGTCTCGTAG |
| MAT | AGTCACATCAAGATCGTTTATGG |
| Alpha | GCACGGAATATGGGACTACTTCG |
| A | ACTCCACTTCAAGTAAGAGTTTG |
| KanB | CTGCAGCGAGGAGCCGTAAT |
| KanC | TGATTTTGATGACGAGCGTAAT |

| ***Diploid yeast strain ID*** | ***Haploid yeast strain ID*** | ***MOBY barcode uptag sequence (5'-3')*** | ***MOBY barcode downtag sequence (3'-5')*** |
| --- | --- | --- | --- |
| d1A2 | h1A2 | GTTCCATACAGCCTAAGTTC | GTTGCATTCACCTACGGTAT |
| d1A3 | h1A3 | CGGCGCTCGATTCTCATTGT | AATCAGACTTGCAGCTTGGG |
| d1A4 | h1A4 | GCCCGCTTTAGTAATATACG | ATAAGCTCTGGGAATAGCCG |
| d1A5 | h1A5 | CGTTAAGCATGGTGTTCAAG | AACTGCGTTGAAGCGTTAAG |
| d1A6 | h1A6 | GAAGTGGTCGTCATACTCTT | TTACTAAGTGGCGAGAAGTC |
| d1A7 | h1A7 | GTAAGCCTTTGTACCCCCCG | TGAGGACCAACATACCTTCA |
| d1A8 |  | AGGAGGAACTTATGTCAAGC | AGATGGCCCTAATTGTCTGC |
| d1A9 |  | GCTAATAACTCTCACGAAGC | AGAGGATACCCACTAAGCGC |
| d1A10 |  | GAGTAGATCACAGAATTTCC | AGTATCACACGAGCTTACAC |
| d1A11 |  | AATGAGGACTCTCGCCAAGC | GGATAATACTAACGACCCGC |
| d1B1 |  | ATCCATAAGGTCGGGCAAGC | GCAGAGTCCTGAACAGCCGC |
| d1B2 |  | CTGGCATAACTCATAAGAGC | TATTAGCACCCGGAAGTCGC |
| d1B3 |  | TTTACCACCATAGACAGAGC | GCATGAGAGCCACCAGTCGC |
| d1B4 |  | GCCCGTTAGAGTAAATAAGC | TAAATAGCCGTGCAACGCGC |
| d1B5 |  | GCGTTACACAGATCATAAGC | AATTAACGACCAGTACGCGC |
| d1B6 |  | TAGTACCCGGAACCTAGGGC | GCATAGGTAACAGGAGTCGC |
| d1B7 |  | CCAGGATGATGAATACGAGC | TTAGGTACACTACACGTCGC |
| d1B8 |  | GAAGCACTTTCAAGCCGAGC | GCTCACCAAAGAAGCGTCGC |
| d1B9 |  | TGTCACGACCACAAAGGAGC | ATTAAGCGCACAGAGGTCGC |
| d1B10 |  | TGACTAAGCCCACCTGGAGC | TACCACTGGGAAGGATTCGC |
| d1B11 |  | TACGCGAACCCATTGGTAGC | GCTGCACATAGAGCACAGGC |
| d1B12 |  | TTACAGCAGCGCAGCTTAGC | TAAATCCCTTTCAAGCAGGC |
| d1C1 |  | TCGACGTGCATCCAGGAATG | GCATTGACGCCAAGCCACTA |
| d1C2 |  | ACATCCTTGGCATGGGAGAT | GCAGATAAGCGCATCGAAGC |
| d1C3 |  | AGACTGCACACATGGTTTCCC | GCCTGAGATTCATGTAGAT |
| d1C4 |  | TCTCGAATGAGTGGCAAAGC | AGCTCAGGACGATTGTAGAT |
| d1C5 |  | ATTGCATCTGAACGCCTGGC | ATCTATGTGGCAGTCGTGCT |
| d1C6 |  | TCATTCGACACATGGCTGGC | CTTATCCTGTAGAGGGTGCT |
| d1C7 |  | TTATGTACCATCAAGCCCAG | TCGCGCATAAGACCAATCGA |
| d1C8 |  | TCTATTATGAGAGCGCCGTG | CAGTCTAATGAATCGGAAGC |
| d1C9 |  | CCTTCTTAGGATGTGAGATC | ATCCATTGAATCCATGTGGC |
| d1C10 |  | GAGACGCCTCGATTACTATG | GCTGTTCGTCCATGCGACCT |
| d1C11 |  | GATACTGTCACACCGTTGCG | GCGCGCCCTATACTATCTAT |
| d1C12 |  | GCCTAGTAGATACGATCAGG | GGCTGATATACGTTCACACT |
| d1D1 |  | GAGATGTGGATCTCACCCAT | GAGCGAGGTCTATATCTTGT |
| d1D2 |  | GACTACCGAGTGCTAGGCAT | GCGAGCCTCCTTCATGTTGT |
| d1D3 |  | GATCAGCCTTTACATTCCTG | GATGGTCTTCCACCTGTGCT |
| d1D4 |  | GACTATGTATTAAGCGTGCG | GGCTACTTCAGCGTT |
| d1D5 |  | GACTAGCGTCCATCTCGTTG | GGTGAGCCTTCATCTCTCGT |
| d1D6 |  | GGCCTGAGAGTCATTTGAAT | GACCTTGAAGTCAGCTATGT |
| d1D7 |  | CTGCGCGGAGATAACAATTA | CTCCGGCATTAGAACATAAC |
| d1D8 |  | TACGAACACCGTCGCAATTA | GGCGGCTAGAGAACCATAAC |
| d1D9 |  | ATCAGGGCCGCAAGTCTGTA | GCGGCAGAACTAATATCAAC |
| d1D10 |  | AGAGGAACCCACGGCCTTTA | CATAGTAGAACAGTTGGCAC |
| d1D11 |  | AGACACGACACACTGGCTTA | GCGGGACACATTGAACACAC |
| d1D12 |  | CGCAGTCAACTTACACGTTA | GGGATTCACACATTAACCAC |
| d1E1 |  | CCATCTGCAATACGTCGTTA | GTCCAGATGACAGAAGCCAC |
| d1E2 |  | GGAGTTCTTCCAATCCAAAC | GGCCGCTAACTAATAATCAC |
| d1E3 |  | CACGAGAACCACGATCTTTA | GGTACACGCGCAGAATGCAC |
| d1E4 |  | CCGACACAGAGAAGTCTTTA | GCACTATTGGCAACATGCAC |
| d1E5 |  | GGCTAAGAACTTCGTACAAT | GGGAAGATACCCTCAACGAC |
| d1E6 |  | TACCGAACAGTGGTTAAAC | GCGCTAACATTACTCAAGAC |
| d1E7 |  | GGATACCATACACATTGGAC | TTGAAGACATCCTACGAACC |
| d1E8 |  | TGGAATACCCACGAACGGAC | GCATAAGACGTTGTGAAACC |
| d1E9 |  | GCACGAGTTAGAACACGGAC | TGAACCACAGTGTTGAAACC |
| d1E10 |  | CTACAGTTGAGAACAGGGAC | AGGAGCACTATATCTCCACC |
| d1E11 |  | TGAGGAACACGGTGCAATAC | AGATGTACGTCAATTCCACC |
| d1E12 |  | GGCCCGAGTACGAATAATAC | TGAGGAAGACCATCTGCACC |
| d1F1 |  | AGAGCTTCAACATTGCTGAC | CACACGGGTAGAAGTGAACC |
| d1F2 |  | TCAAGCGAATCCCATGTGAC | GCGGACATAGCATATTAACC |
| d1F3 |  | ACACGCGAATAAGTCTTGAC | GCGACAGCATGATGAACACC |
| d1F4 |  | CGAGTTACAACAGTCTCTGA | TGCCGAGCTGAACAACGACA |
| d1F5 |  | TAGAACCAGTGTTTAACGGG | ATGAGCTTATCCACGCCTGG |
| d1F6 |  | AGTGCTCTCAGTTCTCCAGT | AGTAGTGCCAGCCTAACATG |
| d1F7 |  | AGGAATCCATCTCAACTGGC | CCGCGTGATATAGATACATG |
| d1F8 |  | AGAGTAGCGACCACTATCTC | GCCGCATAGATATTGGAAAC |
| d1F9 |  | TTGCGGCATGTGACACTCTG | GGCTCATCCTCAAGATTAAC |
| d1F10 |  | ACATAGTCTACAGGATCACG | TTGCCGTGCATCCGATAGCT |
| d1F11 |  | CTTCCCGACAGAGCAAGAGA | GGGCTCAATACACTCGATTA |
| d1F12 |  | GAAGCGTTGAGAATGCCTCC | GTTGACGCACTCACAGTTTC |
| d1G1 |  | GCAGGATACCTAATGATTCC | GCGCAGTAATGCTTTAGAAG |
| d1G2 |  | GGCCACTTCTAACATATAGC | GACACTGGCCTAACTTGCAG |
| d1G3 |  | GCAGTCCTACCAATCTATGA | GCACTCGTGAAATGCACTCA |
| d1G4 |  | CGGAGGCTCAGATGTACTAT | CTGAGCATTCGCCATGTATG |
| d1G5 |  | TTAGCACTCCGACAGCGTAG | TACTCATCGGTAGCCAGTGG |
| d1G6 |  | TATGCGAGTCCAGCTCACTG | GCATTGTCTGAAGCTGTGAG |
| d1G7 |  | TGCACGAGTCTCCAGTGTTG | ATTGGTCATACAGCGCCAGG |
| d1G8 |  | CGCATTGTGACAGATGTTCG | TTGCATCACGCATGTGGTGC |
| d1G9 |  | ACGCTGCTTACAGTGTGTTG | ATGTTGCCCATAGCCCCAGG |
| d1G10 |  | GTATGCTCATCATCCCGTAT | CTATCGAGGATCACGGACTG |
| d1G11 |  | CACTGTGTAGCAGCCGGTAT | ATCTAGGCGGCATGTATTTG |
| d1G12 |  | CCGCGTATAGAATTGTCTGA | GCCACACCTAAATGTAGTCA |
| d1H2 | h1H2 | TCAGGAGACTGCACTACAGG | GCTCTTTAACCATGATGTGC |
| d1H3 | h1H3 | ATAGTGTCACCACAGGTCGC | GCGTATAGCGACGGCCTCTC |
| d1H4 | h1H4 | CCTAGAGGCTAATTTGCATC | CGTGGCTATTCATACATTTC |
| d1H5 | h1H5 | TAGAGAGCTACCACATCATC | AGCATCGCGCTAACTATTTC |
| d1H6 | h1H6 | ACGCACAGGTTATGAATCTC | TGATGACTGACCAGCCTCTG |
| d1H7 | h1H7 | AGGAGCCTATAACTCATCTC | CTAGTGCATAGACTCCTCTG |
| d1H8 |  | CAGAGACTACTACATCGTTC | GCTGGGAGCATGATCTGCAT |
| d1H9 |  | AGGCATCTCACATACTGTTC | TTGTCGCGCAGCTAGTGCAT |
| d1H10 | h1H10 | ATGAGCCCGTGACTGCGATT | GCCGAAGTTATCGCAATGTA |
| d1H11 |  | GGCATACCGACAACGATTCA | ACATCATACGCATCGAGCGG |
