## Supplementary material for "High-throughput analysis of adaptation using barcoded strains of *Saccharomyces cerevisiae*": Supplmental Table 2

| ***Treatment ID*** | ***Evo. Wells*** | ***Barcodes per Evo. Well*** | ***Ploidy*** | ***Evo. Medium*** | ***Evo. Transfer Dilution*** |
| --- | --- | --- | --- | --- | --- |
| 1 | 21 | 2 | Diploid | CM | 1/1000 |
| 2 | 11 | 2 | Haploid | CM | 1/1000 |
| 3 | 11 | 2 | Diploid | CM | 1/250 |
| 4 | 11 | 2 | Diploid | CM | 1/4000 |
| 5 | 11 | 2 | Diploid | CM + 8% EtOH | 1/1000 |
| 6 | 11 | 2 | Diploid | CM + 0.342M NaCl | 1/1000 |
