## Supplementary material for "High-throughput analysis of adaptation using barcoded strains of *Saccharomyces cerevisiae*": Supplmental Table 3

| ***Oligo name*** | ***Full sequence (5'-3')*** | ***Barcode sequence (5'3')*** |
| --- | --- | --- |
| Ion_17 | CCATCTCATCCCTGCGTGTCTCCGACTCAGTCTATTCGTCGATGTCCACGAGGTCTCT | TCTATTCGTC |
| Ion_18 | CCATCTCATCCCTGCGTGTCTCCGACTCAGAGGCAATTGCGATGTCCACGAGGTCTCT | AGGCAATTGC |
| Ion_19 | CCATCTCATCCCTGCGTGTCTCCGACTCAGTTAGTCGGACGATGTCCACGAGGTCTCT | TTAGTCGGAC |
| Ion_20 | CCATCTCATCCCTGCGTGTCTCCGACTCAGCAGATCCATCGATGTCCACGAGGTCTCT | CAGATCCATC |
| Ion_21 | CCATCTCATCCCTGCGTGTCTCCGACTCAGTCGCAATTACGATGTCCACGAGGTCTCT | TCGCAATTAC |
| Ion_22 | CCATCTCATCCCTGCGTGTCTCCGACTCAGTTCGAGACGCGATGTCCACGAGGTCTCT | TTCGAGACGC |
| Ion_23 | CCATCTCATCCCTGCGTGTCTCCGACTCAGTGCCACGAACGATGTCCACGAGGTCTCT | TGCCACGAAC |
| Ion_24 | CCATCTCATCCCTGCGTGTCTCCGACTCAGAACCTCATTCGATGTCCACGAGGTCTCT | AACCTCATTC |
| Ion_25 | CCATCTCATCCCTGCGTGTCTCCGACTCAGCCTGAGATACGATGTCCACGAGGTCTCT | CCTGAGATAC |
| Ion_26 | CCATCTCATCCCTGCGTGTCTCCGACTCAGTTACAACCTCGATGTCCACGAGGTCTCT | TTACAACCTC |
| Ion_27 | CCATCTCATCCCTGCGTGTCTCCGACTCAGAACCATCCGCGATGTCCACGAGGTCTCT | AACCATCCGC |
| Ion_28 | CCATCTCATCCCTGCGTGTCTCCGACTCAGATCCGGAATCGATGTCCACGAGGTCTCT | ATCCGGAATC |
| Ion_29 | CCATCTCATCCCTGCGTGTCTCCGACTCAGTCGACCACTCGATGTCCACGAGGTCTCT | TCGACCACTC |
| Ion_30 | CCATCTCATCCCTGCGTGTCTCCGACTCAGCGAGGTTATCGATGTCCACGAGGTCTCT | CGAGGTTATC |
| Ion_31 | CCATCTCATCCCTGCGTGTCTCCGACTCAGTCCAAGCTGCGATGTCCACGAGGTCTCT | TCCAAGCTGC |
| Ion_32 | CCATCTCATCCCTGCGTGTCTCCGACTCAGTCTTACACACGATGTCCACGAGGTCTCT | TCTTACACAC |
| Ion_33 | CCATCTCATCCCTGCGTGTCTCCGACTCAGTTCTCATTGAACGATGTCCACGAGGTCTCT | TTCTCATTGAAC |
| Ion_34 | CCATCTCATCCCTGCGTGTCTCCGACTCAGTCGCATCGTTCGATGTCCACGAGGTCTCT | TCGCATCGTTC |
| Ion_35 | CCATCTCATCCCTGCGTGTCTCCGACTCAGTAAGCCATTGTCGATGTCCACGAGGTCTCT | TAAGCCATTGTC |
| Ion_36 | CCATCTCATCCCTGCGTGTCTCCGACTCAGAAGGAATCGTCGATGTCCACGAGGTCTCT | AAGGAATCGTC |
| Ion_37 | CCATCTCATCCCTGCGTGTCTCCGACTCAGCTTGAGAATGTCGATGTCCACGAGGTCTCT | CTTGAGAATGTC |
| Ion_38 | CCATCTCATCCCTGCGTGTCTCCGACTCAGTGGAGGACGGACGATGTCCACGAGGTCTCT | TGGAGGACGGAC |
| Ion_39 | CCATCTCATCCCTGCGTGTCTCCGACTCAGTAACAATCGGCGATGTCCACGAGGTCTCT | TAACAATCGGC |
| Ion_40 | CCATCTCATCCCTGCGTGTCTCCGACTCAGCTGACATAATCGATGTCCACGAGGTCTCT | CTGACATAATC |
| Ion_R1 | CCTCTCTATGGGCAGTCGGTGATAGCGCTTAGGATGTCGACCTGCAGCGTACG | TCGCGAATC |
| Ion_R2 | CCTCTCTATGGGCAGTCGGTGATGACTGATACGATGTCGACCTGCAGCGTACG | CTGACTATG |
| Ion_R3 | CCTCTCTATGGGCAGTCGGTGATATTCAATTCGATGTCGACCTGCAGCGTACG | TAAGTTAAG |
| Ion_R4 | CCTCTCTATGGGCAGTCGGTGATCTGAAACCGGATGTCGACCTGCAGCGTACG | GACTTTGGC |
| Ion_R5 | CCTCTCTATGGGCAGTCGGTGATGTTGGGCCGGATGTCGACCTGCAGCGTACG | CAACCCGGC |
| Ion_R6 | CCTCTCTATGGGCAGTCGGTGATTGGTATGCCGATGTCGACCTGCAGCGTACG | ACCATACGG |
| Ion_R7 | CCTCTCTATGGGCAGTCGGTGATCAAGCGAGCGATGTCGACCTGCAGCGTACG | GTTCGCTCG |
| Ion_R8 | CCTCTCTATGGGCAGTCGGTGATTCCCGACCGGATGTCGACCTGCAGCGTACG | AGGGCTGGC |
| Ion_R9 | CCTCTCTATGGGCAGTCGGTGATGAGGATGATGATGTCGACCTGCAGCGTACG | CTCCTACTA |
| Ion_R10 | CCTCTCTATGGGCAGTCGGTGATAAATCGAATGATGTCGACCTGCAGCGTACG | TTTAGCTTA |
| Ion_R11 | CCTCTCTATGGGCAGTCGGTGATAGCTGCCGAGATGTCGACCTGCAGCGTACG | TCGACGGCT |
| Ion_R12 | CCTCTCTATGGGCAGTCGGTGATAGAGGCTGCGATGTCGACCTGCAGCGTACG | TCTCCGACG |
| Ion_R13 | CCTCTCTATGGGCAGTCGGTGATAACGTGAGGGATGTCGACCTGCAGCGTACG | TTGCACTCC |
